# The two enantiomers of 2-hydroxyglutarate differentially regulate cytotoxic T cell function

**DOI:** 10.1101/2022.08.18.504372

**Authors:** Iosifina P. Foskolou, Pedro P Cunha, Eleanor Minogue, Benoit P Nicolet, Aurelie Guislain, Christian Jorgensen, Nordin D Zandhuis, Laura Barbieri, David Bargiela, Demitris Nathanael, Petros Tyrakis, Asis Palazon, Monika C Wolkers, Randall S Johnson

## Abstract

2-hydroxyglutarate (2HG) is a by-product of the TCA cycle, and is readily detected in the tissues of healthy individuals. 2HG is found in two enantiomeric forms: S-2HG and R-2HG. Here, we investigate the differential roles of these two enantiomers in CD8+ T cell biology, where we found they had highly divergent effects on proliferation, differentiation, and T cell function. We show here an analysis of structural determinants that likely underlie these differential effects on specific a-ketoglutarate (aKG)-dependent enzymes. Treatment of CD8+ T cells with exogenous S-2HG, but not R-2HG, increased CD8+ T cell fitness *in vivo,* and enhanced anti-tumour activity. These data show that S-2HG and R-2HG should be considered as two distinct and important actors in the regulation of T cell function.

## INTRODUCTION

There are three known metabolites with structural similarities to alpha-ketoglutarate (aKG) that have been shown to inhibit a range of aKG-dependent enzymes. Two of these, succinate and fumarate, are tricarboxylic acid (TCA) cycle metabolites with essential metabolic roles, and the third is 2-hydroxyglutarate (2HG), a physiological by-product of the TCA cycle. 2HG is found in two forms: the S-form (also known as L-2HG) and the R-form (also known as D-2HG). Intracellular accumulation of 2HG is derived by the reduction of aKG to either R-2HG, or to S-2HG (Du and Hu, 2021). Both enantiomers of 2HG can be detected in most body fluids of healthy individuals, and their concentrations can reach near millimolar levels in urine and serum (Fitzpatrick et al., 2020; Strain et al., 2020). Although 2HG likely has numerous physiological roles, it has been primarily studied in the context of specific diseases, such as 2-hydroxyglutaric acidurias (2HGAs). 2HGAs are rare diseases characterised by accumulation of 2HG (S-2HG and/or R-2HG) in body fluids, and result in a series of psychiatric and neurological symptoms (Kranendijk et al., 2012; Rodrigues et al., 2017).

R-2HG levels are increased in tumours harbouring mutations in isocitrate dehydrogenase (IDH1/2mut), which include glioblastomas, acute myeloid leukaemias (AML), chondrosarcomas, osteosarcomas, cholangiocarcinomas, and prostate cancers (Du and Hu, 2021; Mardis et al., 2009; Parsons et al., 2008; Yan et al., 2009). Accumulation of S-2HG has been reported in clear cell renal cell carcinomas (ccRCC) and pancreatic cancers (Gupta et al., 2021; Shim et al., 2014). However, 2HG’s role in cancer initiation, progression and drug response is more complex than initially thought. For example, IDHmut glioma and AML patients tend to have better overall survival rates than IDH wild type (IDHwt) patients (Chou et al., 2011; Eckel-Passow et al., 2015; Parsons et al., 2008; Patel et al., 2012; Yan et al., 2009). In addition, in several cancer models, both S-2HG and R-2HG have been shown to have the capacity to reduce tumour growth when they are exogenously administrated or endogenously produced (Fu et al., 2015; Su et al., 2018).

Due to the differential 2HG accumulation in the tumour microenvironment, it is becoming increasingly important to determine whether the two enantiomers of 2HG act in a similar fashion in immune cells, or whether the two enantiomers should be considered as clearly separate biological agents from a mechanistic standpoint. As both enantiomers can affect cytotoxic T cell differentiation, proliferation, and function (Du and Hu, 2021; Foskolou et al., 2020; Tyrakis et al., 2016; Yang et al., 2022), we wished to determine the extent to which they have clearly differentiable roles in CD8+ T cell function. We show here that the two enantiomers are qualitatively separable in their actions and should be considered as two separate actors in regulating T cell function.

## RESULTS

### Human CD8+ T cells express different surface markers when treated with cell-permeable S-2HG or R-2HG

The structural similarity of both enantiomers of 2HG to aKG causes them to act as competitive inhibitors of aKG-dependent enzymes (Fig. 1A) (Chowdhury et al., 2011; Xu et al., 2011). To investigate the differential effects of the two enantiomers of 2HG on human CD8+ T cells, we used cell-permeable octyl ester forms of S-2HG (OE-S-2HG) and R-2HG (OE-R-2HG). To minimise donor-to-donor variation, naïve CD8+ T cells from healthy individuals were used. Neither OE-S-2HG nor OE-R-2HG had an effect on CD8+ T cell viability when used in physiologically relevant concentrations (0.4 mM) (Fitzpatrick et al., 2020; Tyrakis et al., 2016) for up to 12 days of culture, under culture conditions of OE-S-2HG or OE-R-2HG supplementation every two days (Fig. 1B).

**Figure 1:**
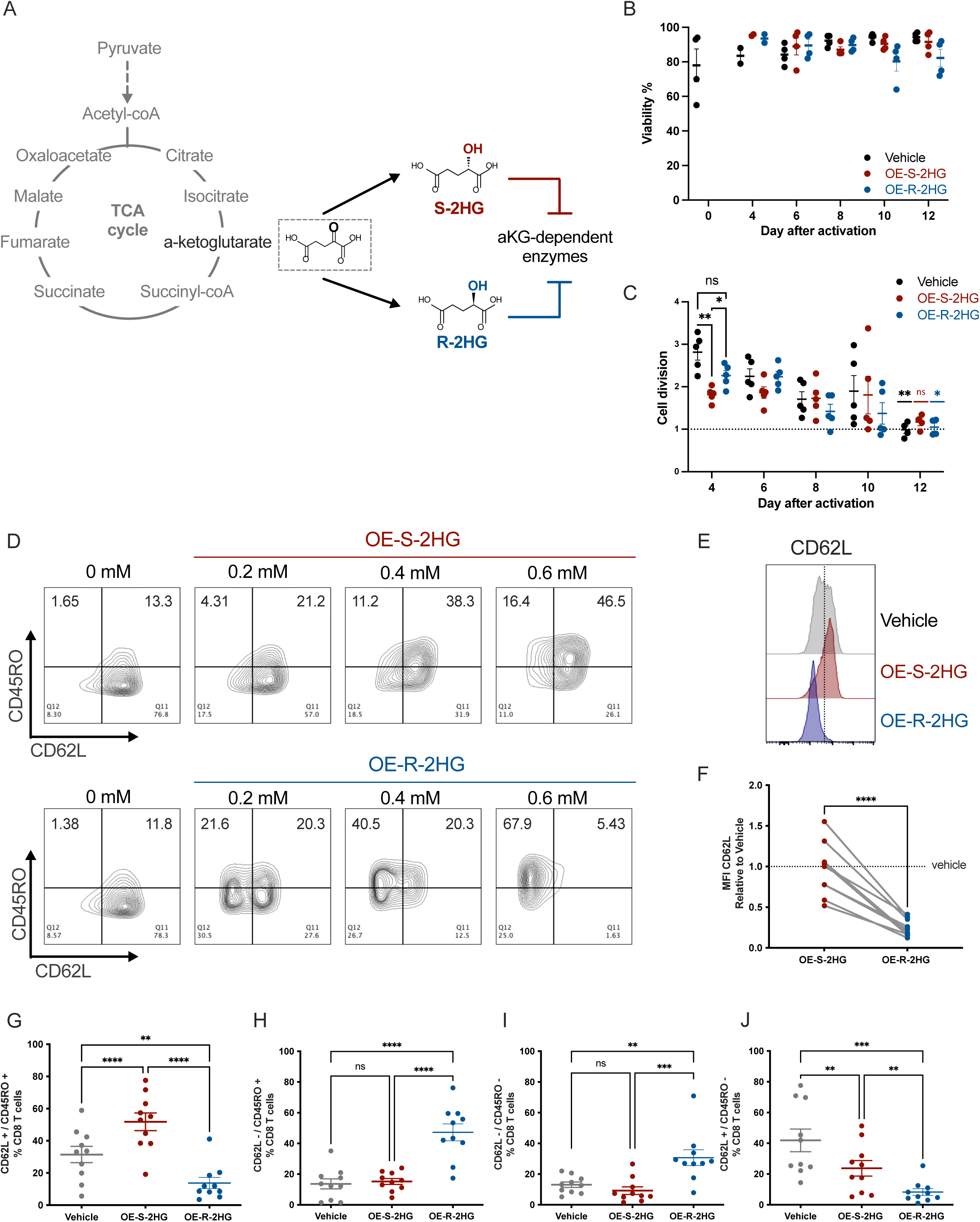
Human CD8+ T cells express different surface markers when treated with OE-S-2HG or OE-R-2HG. **(A)** Schematic representation of the TCA cycle and chemical structures of a-ketoglutarate (aKG), S-2hydroxyglutarate (S-2HG) and R-2hydroxyglutarate (R-2HG). S-2HG and R-2HG can act as competitive inhibitors of aKG-dependent enzymes, due to their structural similarity with aKG. **(B)** Viability of CD8+ T cells treated with Octyl-ester-S-2HG (OE-S-2HG; 0.4 mM), Octyl-ester-R-2HG (OE-R-2HG; 0.4 mM) or vehicle (H_2_O) for the indicated days. Viability was determined by an automated cell counter. Data are represented as mean ± SEM. **(C)** Cell division of CD8+ T cells treated with OE-S-2HG (0.4 mM), OE-R-2HG (0.4 mM) or vehicle (H_2_O) for the indicated days. Mixed-effects analysis with Tukey’s multiple comparison test was used. Data are represented as mean ± SEM. **(D)** Flow cytometry plots of CD8+ T cells showing surface expression of CD62L and CD45RO. Cells were treated with vehicle (H_2_O) or increasing concentrations of OE-S-2HG or OE-R-2HG and analysed at day 12 by flow cytometry. Representative plots of n= 3 is shown. **(E)** Histograms of representative flow cytometry plots for CD62L expression on CD8+ T cells treated with OE-S-2HG (0.4 mM), OE-R-2HG (0.4 mM) or vehicle (H_2_O) for 12 days. **(F)** Fold change of median fluorescence intensity (MFI) of CD62L for CD8+ T cells treated with OE-S-2HG (0.4 mM) or OE-R-2HG (0.4 mM) relative to vehicle (H_2_O). Cells were analysed at day 12 by flow cytometry. Each data point represents a donor (n= 9; from 6 independent experiments). Unpaired two-tailed Student t test was used. **(G-J)** Cells were treated with OE-S-2HG (0.4 mM), OE-R-2HG (0.4 mM) or vehicle (H_2_O) and the proportion of (G) CD62L+/CD45RO+; (H) CD62L-/CD45RO+; (I) CD62L-/CD45RO-; (J) CD62L+/CD45RO-cells is shown (%CD8+ T cells). Cells were analysed at day 12 by flow cytometry. Each data point represents a donor (n= 10; from 7 independent experiments). Data are represented as mean ± SEM. RM one-way ANOVA with Tukey’s multiple comparison test was used. For all panels naïve CD8+ T cells were isolated and activated with CD3/CD28 beads and cultured with IL2 (30 U/mL) in the presence of OE-S-2HG (0.4 mM), OE-R-2HG (0.4 mM) or vehicle (H_2_O) from day 0 to 12, unless otherwise stated.

Early after activation, CD8+ T cells treated with OE-S-2HG proliferated less (Supp. Fig. 1A) and had a slower rate of cell division (day 4; Fig. 1C) when compared to CD8+ T cells treated with OE-R-2HG or vehicle. However, cell division of OE-S-2HG treated CD8+ T cells remained stable during later time points, whereas cell division of vehicle and OE-R-2HG treated CD8+ T cells was gradually and significantly decreased (Fig. 1C). On day 12, OE-S-2HG treated CD8+ T cells had a cell division rate of (1.169±0.17), followed by OE-R-2HG (1.049±0.19) and vehicle (0.9879±0.18) (Fig. 1C). These data cumulatively indicate that OE-S-2HG treated CD8+ T cells proliferate slower after activation, and that their proliferation is less decreased at later timepoints relative to OE-R-2HG and vehicle treated cells.

We then examined the effect of both enantiomers on the expression of several surface markers related to human CD8+ T cell differentiation. Increasing concentrations of OE-S-2HG led to a dose-dependent increase of CD62L+/CD45RO+ (Fig. 1D) and CCR7+/CD45RO+ (Supp. Fig. 1B) CD8+ T cells, indicating an increase in the markers associated with early differentiated memory-like CD8+ T cells. Although increasing concentrations of OE-R-2HG increased the CCR7+/CD45RO+ population (Supp. Fig. 1B), they led to a dose-dependent loss of CD62L and an increase in CD62L-/CD45RO+ (Fig. 1D-F) CD8+ T cells. The differential expression of CD62L and CCR7 determines the differential migratory tendencies of CD8+ T cells. T cells expressing both CD62L and CCR7 can recirculate between lymphoid tissues and peripheral blood (Sallusto et al., 1999), whereas cells that express CCR7 but are CD62L negative are usually T cells exiting peripheral tissues, and are found in afferent lymphatics (Debes et al., 2005; Mackay et al., 1992).

At 0.4 mM, both OE-S-2HG and OE-R-2HG significantly increased the CCR7+/CD45RO+ population compared to vehicle (Supp. Fig. 1C), but only OE-S-2HG increased the CD62L+/CD45RO+ population compared to both vehicle and OE-R-2HG (Fig. 1G). The OE-R-2HG enantiomer significantly increased the CD62L-/CD45RO+ and CD62L-/CD45RO-populations, and decreased the CD62L+/CD45RO-populations, relative to treatment with either vehicle or OE-S-2HG (Fig. 1H-J). Neither OE-S-2HG nor OE-R-2HG had an effect on CCR7-/CD45RO+ and CCR7-/CD45RO-populations, and only OE-S-2HG treated populations had decreased CCR7+/CD45RO-CD8+ T cells, when compared to vehicle treated cells (Supp. Fig. 1D-F).

Interestingly, OE-R-2HG (0.4 mM) treated CD8+ T cells increased expression of the homing marker CCR7, but showed decreased expression of the co-stimulatory marker CD28, relative to CD8+ T cells treated with OE-S-2HG (Supp. Fig. 1G-H). OE-R-2HG treated cells showed a moderate but not significant increase of the activation/checkpoint marker PD-1, and the transcription factor TOX (Supp. Fig. 1I-J). In summary, CD8+ T cells treated with the two cell permeable forms of 2HG show significant differences in cell division, proliferation, and expression of multiple surface markers.

### Transcriptome analysis reveals distinct differences between OE-S-2HG and OE-R-2HG treated human CD8+ T cells

To assess if the observed differences in cell proliferation and surface markers were due to transcriptome-related changes between OE-S-2HG and OE-R-2HG treated CD8+ T cells, we performed RNA-Seq analysis of naïve CD8+ T cells early (day 5) and late (day 12) after activation. Cells were isolated, activated and treated every two days with OE-S-2HG (0.4 mM), OE-R-2HG (0.4 mM) or vehicle (H_2_O). Hierarchical clustering revealed distinct clusters of transcript expression depending on treatment with OE-S-2HG and OE-R-2HG, or vehicle, at both day 5 (Supp. Fig. 2A) and day 12 (Fig. 2A). Volcano plot analysis identified a total of 361 differentially expressed genes (log_2_ fold change > 0.5; *p.adj* < 0.05) between OE-S-2HG and OE-R-2HG treated cells on day 12 of treatment (Fig. 2B). Hierarchical clustering of significant genes revealed differential expression of transcription factors in OE-S-2HG and OE-R-2HG treated cells (Fig. 2C), as well as cluster of differentiation (CD) molecules and secreted molecules (Supp. Fig. 2B-C). Interestingly, in concordance with protein measurements (Fig. 1E-F and Supp. Fig. 1H), OE-S-2HG treated CD8+ T cells expressed higher *SELL* (CD62L) and *CD28* transcript levels compared to OE-R-2HG treated cells (Fig. 2B). Furthermore, OE-S-2HG treated cells showed higher transcript levels of some genes that are preferentially expressed in CD8+ naïve (T_N_) and/or central memory (T_CM_) T cells, including *SELL* (CD62L), *CD28*, *GPR15*, *NT5E* (CD73), *FUT7* and *ZNF69* (Table 1, Fig. 2B).

**Figure 2:**
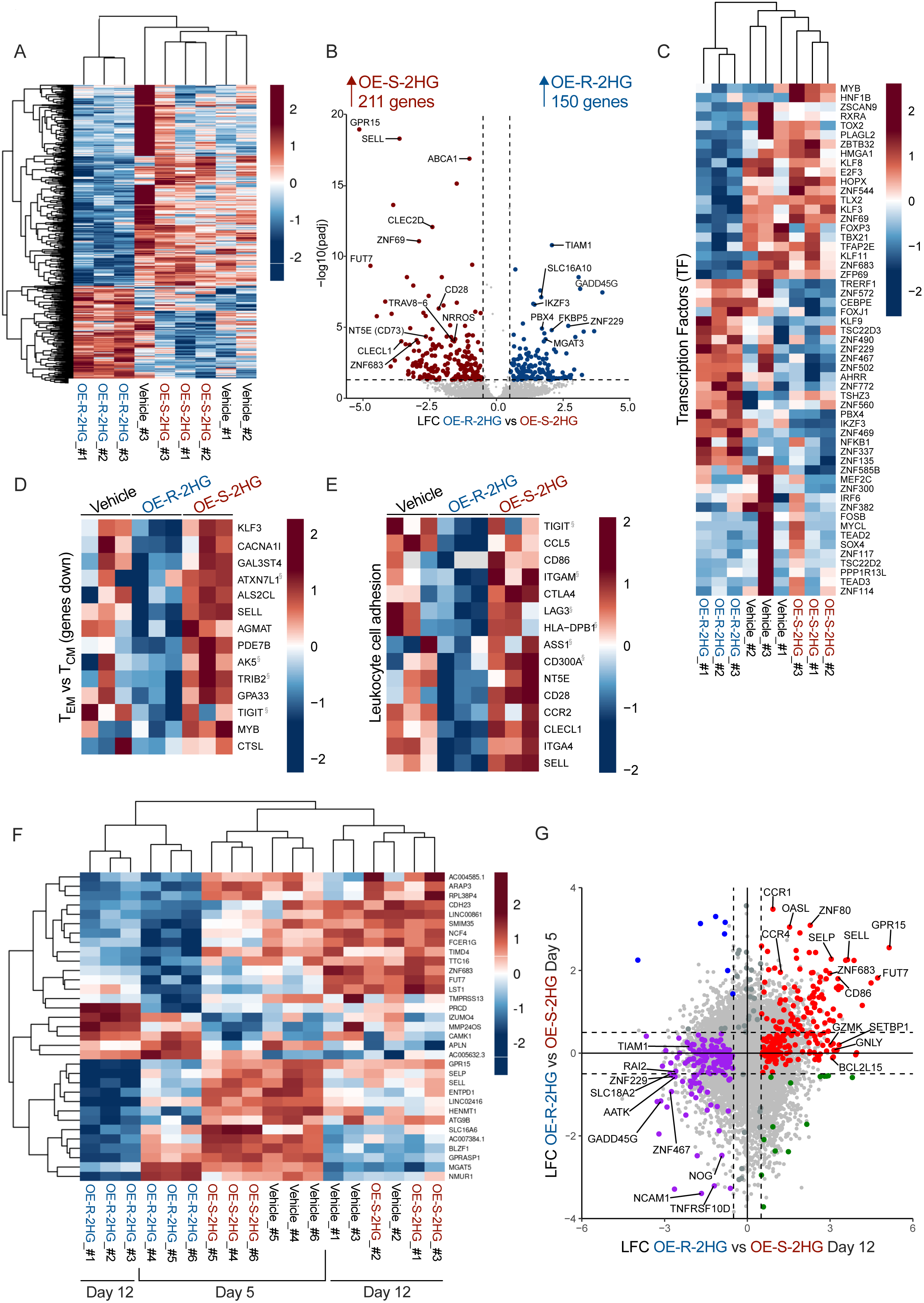
OE-S-2HG and OE-R-2HG treated human CD8+ T cells have distinct transcriptomes. Naïve CD8+ T cells were isolated from 6 individual donors over 3 independent experiments, activated and treated with OE-S-2HG (0.4 mM), OE-R-2HG (0.4 mM) or vehicle (H_2_O). The cells were collected either on day 5 (3 donors) or on day 12 (3 donors) and RNA-Seq analysis followed. **(A)** Heatmap of hierarchically clustered genes in CD8+ T cells treated with OE-S-2HG (0.4 mM), OE-R-2HG (0.4 mM) or vehicle (H_2_O) at day 12 of culture. **(B)** Volcano plot showing log_2_ fold change (LFC; x-axis) and -log_10_ *p-adjusted* value (y-axis) of transcripts differentially expressed in OE-R-2HG-treated CD8+ T cells *vs* OE-S-2HG-treated CD8+ T cells. Samples from day 12 of treatment are shown. Coloured dots represent log_2_ fold change >0.5 and *p.adj*<0.05. **(C)** Heatmap of hierarchically clustered genes in CD8+ T cells treated with OE-S-2HG (0.4 mM), OE-R-2HG (0.4 mM) or vehicle (H_2_O) at day 12 of culture. Statistically significant differentially expressed hits of transcription factors are shown. **(D-E)** Heatmaps of standardized gene expression (Z score) in treated CD8+ T cells: downregulated genes in T_EM_ cells compared to T_CM_ cells; (D) genes involved in leukocyte cell adhesion. Gene sets were obtained from ToppGene. Red and blue colours indicate increased and decreased expression respectively. Genes marked with ‘§’ were non-statistically significant hits. Samples from day 12 of treatment are shown. **(F)** Heatmap of hierarchically clustered genes in CD8+ T cells treated with OE-S-2HG (0.4 mM), OE-R-2HG (0.4 mM) or vehicle (H_2_O) both at day 5 and at day 12 of culture. **(G)** Log_2_ fold change (LFC) of OE-R-2HG vs OE-S-2HG day 5 compared to this of day 12. Dotted lines represent absolute LFC of 0.5. The red dots indicate statistically significant genes for OE-S-2HG upregulated at day 12 (LFC > 0.5), which were also either upregulated or unchanged at day 5 (LFC > -0.5). The purple dots indicate statistically significant genes for OE-R-2HG upregulated at day 12 (LFC < - 0.5), which were also either upregulated or unchanged at day 5 (LFC < 0.5).

**Table 1:**
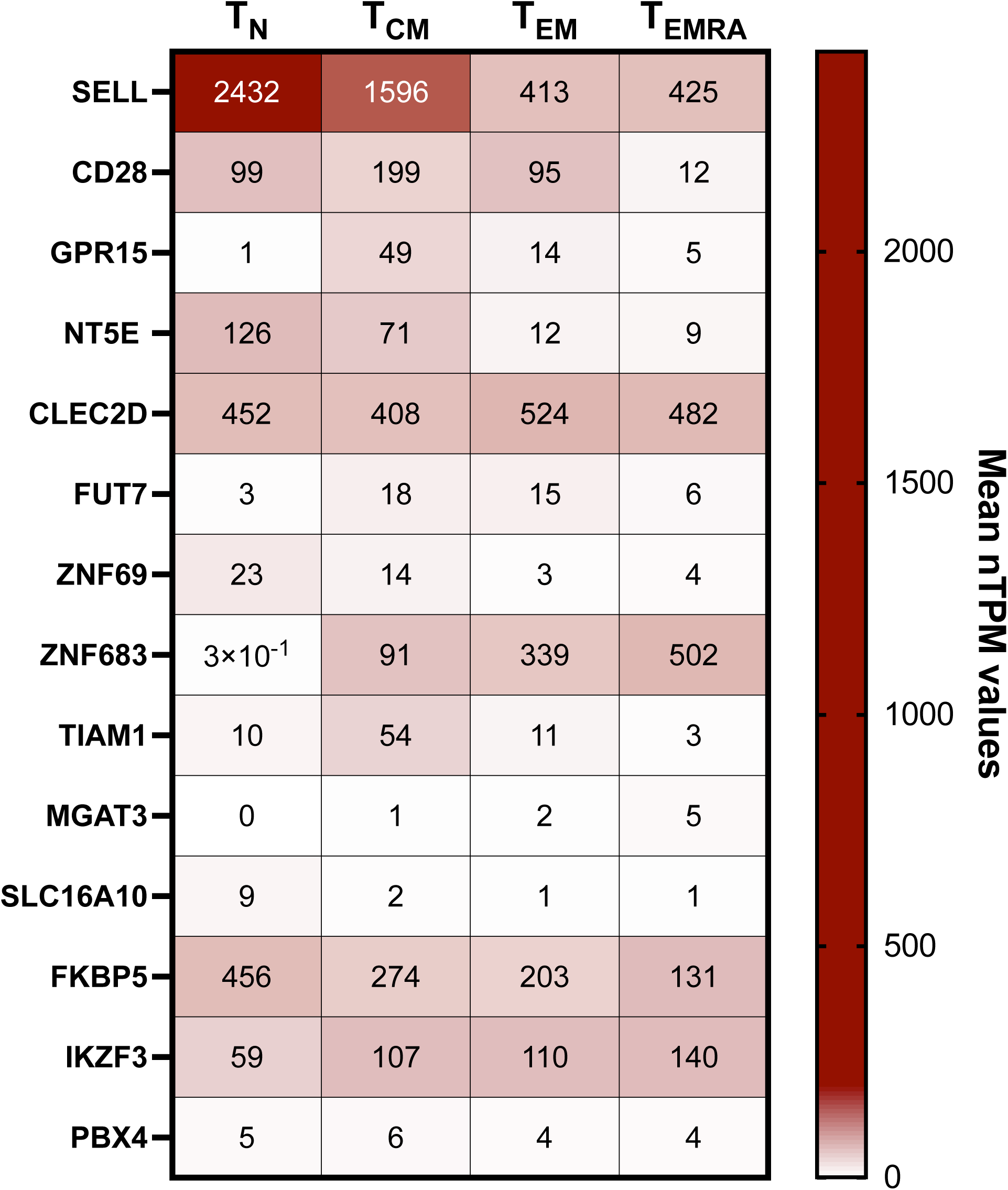
Expression levels of genes upregulated in OE-S-2HG or OE-R-2HG samples in specific CD8+ T cell subsets. Gene expression of specific targets is shown for naïve CD8+ T cells (T_N_), central memory CD8+ T cells (T_CM_), effector memory CD8+ T cells (T_EM_) and terminally differentiated effector memory CD8+ T cells (T_EMRA_). The values of each target were taken by the Monaco dataset (Monaco et al., 2019) through Human Protein Atlas. The data were normalised using trimmed mean of M values (TMM) to allow for between-sample comparisons. The resulting normalised transcript expression values, denoted nTPM, were calculated for each gene in every sample.

By using publicly available datasets, we observed that OE-R-2HG decreased the expression of genes whose expression is usually low in effector memory (T_EM_) CD8+ T cells when compared to T_CM_ cells; with the opposite being the case for OE-S-2HG treated cells (Fig. 2D). Interestingly, the gene signature of OE-R-2HG treated cells was not typical of effector / effector memory cells; as markers associated with cytotoxicity and effector functions were lower in OE-R-2HG treated cells relative to OE-S-2HG and vehicle treated cells (e.g., *ZFN683, HOPX, ZBTB32* and *TBX21*) (Fig. 2B-C). In addition, OE-R-2HG treated cells had higher expression of *IKZF3* compared to OE-S-2HG treated cells (Fig. 2B-C): this gene has been shown to be a key repressor of effector function in CD4+ T cells (Bernardi et al., 2021).

OE-S-2HG treated cells showed increased expression of genes important for leukocyte cell adhesion (Fig. 2E) and T cell activation (Supp. Fig. 2D). Furthermore, gene set enrichment analysis (GSEA) revealed that OE-S-2HG treated T cells displayed enrichment of genes associated with early T lymphocytes, cell cycling genes and E2F3 targets compared to OE-R-2HG (Supp. Fig. 2E-G). E2F3 is a transcription factor which interacts directly with the retinoblastoma protein (pRB) to regulate the expression of genes involved in the cell cycle (Dyson, 1998). These results support our observation that OE-S-2HG treated cells have a moderately higher cell division rate than OE-R-2HG treated cells at day 12 of treatment (Fig. 1C).

Finally, we questioned whether different time points of treatment with the two enantiomers (day 5 and day 12) resulted in different gene expression profiles. Interestingly, hierarchical clustering of the two time points showed close clustering of OE-R-2HG treated CD8+ T cells, indicating a treatment effect (Fig. 2F). In addition, the two most separated T cell populations were these treated for 12 days with OE-R-2HG and OE-S-2HG (Fig. 2F), indicating that the longer the cells are treated with either of the two 2HG compounds, the greater the divergence in their transcriptomes. This finding was supported by the log_2_ fold change (LFC) analysis of OE-R-2HG *vs* OE-S-2HG at day 5 compared to day 12 (Fig. 2G and Supp. Fig. 2H-I). Interestingly, for OE-S-2HG treated cells, genes such as *SELL*, *SELP*, *FUT7*, *GPR15*, *ZNF683, CCR1* and *CCR4* were upregulated at both time points measured (Fig. 2G). These data indicate that OE-S-2HG and OE-R-2HG treated cells have undergone a deep rearrangement of their transcriptomes.

### OE-S-2HG and OE-R-2HG differentially inhibit aKG-dependent enzymes

We next sought to unravel the molecular mechanisms underlying the different transcriptional profiles of OE-S-2HG and OE-R-2HG treated CD8+ T cells. Because of their structural similarities with aKG, both S-2HG and R-2HG can act as competitive inhibitors of aKG-dependent enzymes (Chowdhury et al., 2011; Xu et al., 2011). Some aKG-dependent enzymes are epigenetic modulators, including histone and DNA demethylases (Chowdhury et al., 2011). Therefore, we first assessed the effect of OE-S-2HG and OE-R-2HG treatment on histone modifications in CD8+ T cells.

OE-S-2HG treatment significantly increased histone 3 lysine 9 acetylation (H3K9ac) compared to OE-R-2HG, whereas OE-R-2HG slightly increased H3K9 tri-methylation (H3K9me3), and significantly increased H3K9 di-methylation (H3K9me2) (Fig. 3A-B). In addition, OE-R-2HG treated cells showed substantially increased H3K27 tri-methylation (H3K27me3) levels, at the expense of H3K27 acetylation (H3K27ac) compared to OE-S-2HG treated cells (Fig. 3A,C). These data are further supported by the broad increase of gene expression of histone deacetylases (HDAC) in OE-R-2HG treated T cells, and a decrease thereof in the OE-S-2HG samples (Supp. Fig. 3A). We next checked genes that are regulated by Enhancer of zeste 2 (EZH2) and SUZ12, two members of the polycomb repressive complex 2 (PRC2), which regulates H3K27me deposition (Supp. Fig. 3B) (van Mierlo et al., 2019). EZH2 was found to be essential for effector CD8+ T cell expansion, and it has been shown that PRC2 deficiency impairs effector CD8+ T cell differentiation, but minimally impacts memory CD8+ T cell maturation (Gray et al., 2017). Indeed, SUZ12 target genes were increased in OE-S-2HG treated T cells and decreased in OE-R-2HG treated T cells (Supp. Fig. 3C). Similarly, EZH2-associated genes were significantly increased in OE-R-2HG treated T cells (Supp. Fig. 3D).

**Figure 3:**
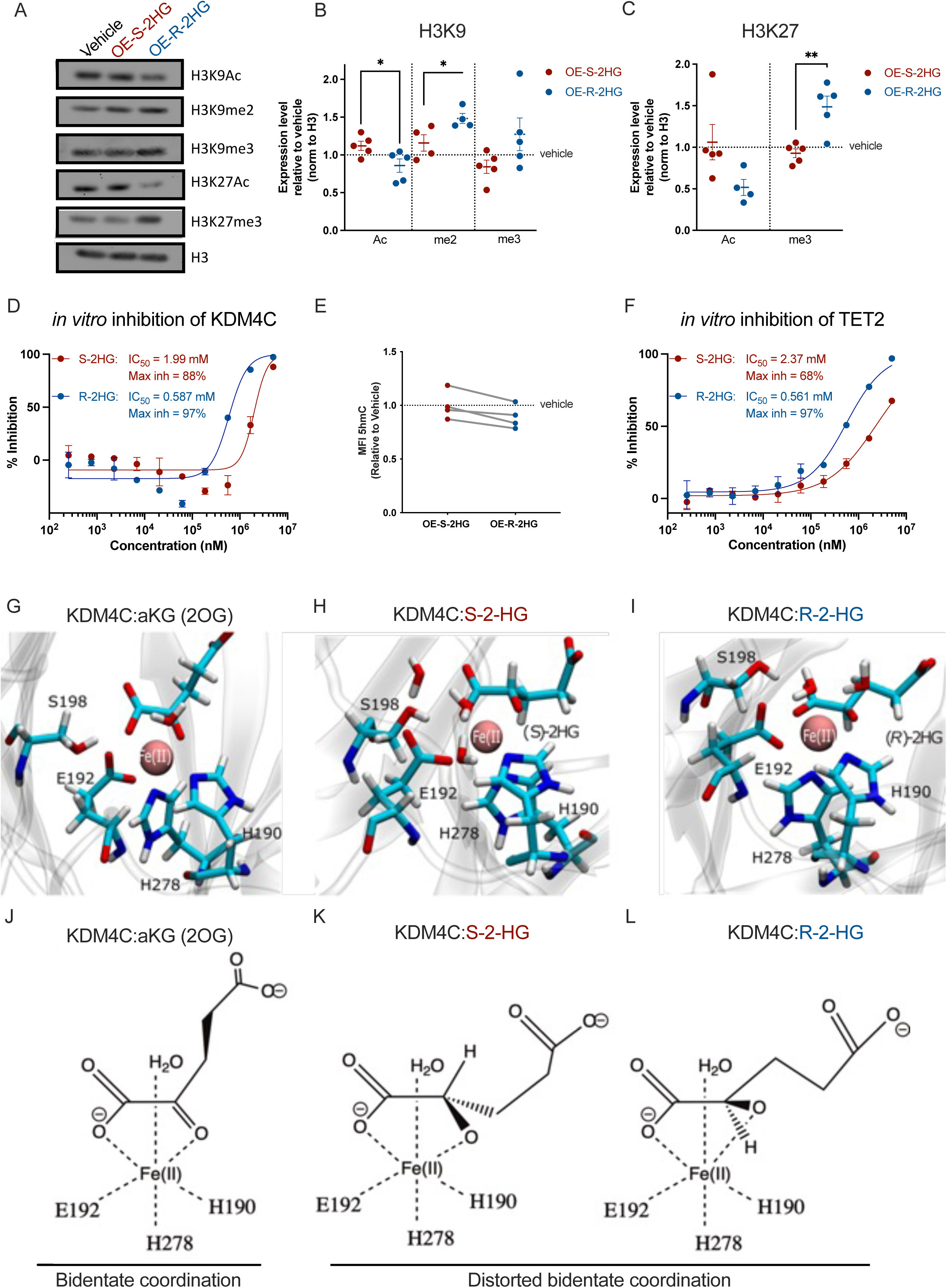
OE-S-2HG and OE-R-2HG have different inhibition potencies for some aKG-dependent enzymes. **(A)** Western Blot analysis of CD8+ T cells treated with OE-S-2HG (0.4 mM), OE-R-2HG (0.4 mM) or vehicle (H_2_O) from day 0 to 12. The cells were lysed at day 12 and different histone marks were assessed. Representative images of n=4 is shown. **(B-C)** Quantification of the expression levels of different histone marks shown in (A) relative to total H3 for (B) H3K9 and (C) H3K27. Each data point represents a donor (n= 4-5; from 4 independent experiments). Data are represented as mean ± SEM. Unpaired two-tailed Student t test was used. **(D)** *In vitro* enzymatic inhibition assay for the KDM4C enzyme. Inhibition was determined by using increasing concentrations of S-2HG or R-2HG. Data are represented as mean ± SD. Fold change of median fluorescence intensity (MFI) of 5hmC for CD8+ T cells treated with OE-S-2HG (0.4 mM) or OE-R-2HG (0.4 mM) relative to vehicle (H_2_O). Cells were analysed at day 7 by flow cytometry. Each data point represents a donor (n= 4; from 2 independent experiments). Unpaired two-tailed Student t test was used. (D) *In vitro* enzymatic inhibition assay for the TET2 enzyme. Inhibition was determined by using increasing concentrations of S-2HG or R-2HG. Data are represented as mean ± SD. **(G-I)** Representative snapshots of the catalytic site of KDM4C protein (PDB id 4XDO; resolution 1.97 Å) in complex with (G) aKG (KDM4C:2OG:Fe(II)), (H) S-2HG (KDM4C:2HG-(S):Fe(II)) and (I) R-2HG (KDM4C:2HG-(R):Fe(II)). **(J)** aKG (KDM4C:2OG:Fe(II)) maintains the pocket integrity from the crystal structure and is found in a bidentate coordination. **(K)** In KDM4C:2HG-(S):Fe(II), the backbone of S-2HG rearranges to maintain a pseudo-bidentate coordination of Fe(II). **(L)** In KDM4C:2HG-(R):Fe(II), the backbone of R-2HG resembles aKG, and no conformational rearrangement occurs. The enzymatic pocket of KDM4C:2HG-(S):Fe(II) and KDM4C:2HG-(R):Fe(II) contains additional waters near the coordination sphere of Fe(II), as a result of the disruption induced by the change from aKG to S-2HG or R-2HG. For panels A-C naïve CD8+ T cells were isolated and activated with CD3/CD28 beads and cultured with IL2 (30 U/mL) in the presence of OE-S-2HG (0.4 mM), OE-R-2HG (0.4 mM) or vehicle (H_2_O) from day 0 to 12. For panel E total CD8+ T cells were isolated and activated with CD3/CD28 beads and cultured with IL2 (30 U/mL) in the presence of OE-S-2HG (0.4 mM), OE-R-2HG (0.4 mM) or vehicle (H_2_O) from day 0 to 7.

To determine whether S-2HG or R-2HG could functionally alter the activity of demethylases, we performed *in vitro* enzymatic activity assays for the H3K9 demethylase KDM4C. The product of the reaction was detected by a highly specific antibody recognising the demethylated substrate, and enzymatic activity measured by immunocytochemistry using fluorescence resonance energy transfer (FRET) technology. In agreement with our cellular data, we observed that R-2HG was a more potent inhibitor than S-2HG, and that the maximum percentage of KDM4C inhibition with our assay was 97% for R-2HG and 88% for S-2HG (Fig. 3D).

Finally, we checked the effect of OE-S-2HG and OE-R-2HG treatment on DNA methylation. DNA methylation directly dictates CD8+ T cell differentiation and function (Correa et al., 2020). The ten-eleven translocation (TET) enzymes are aKG-dependent DNA demethylases that catalyse the oxidation of 5-methylcytosine (5mC) to 5-hydroxymethylcytosine (5hmC) (Matuleviciute et al., 2021; Tahiliani et al., 2009). To determine if OE-S-2HG or OE-R-2HG treatment affects DNA methylation, we treated CD8+ T cells for seven days with OE-S-2HG, OE-R-2HG, or vehicle, and determined total 5hmC levels by flow cytometry. For all donors tested, a moderate decrease of the median fluorescent intensity (MFI) of 5hmC was observed in OE-R-2HG treated CD8+ T cells compared to OE-S-2HG treated cells (Fig. 3E). We performed *in vitro* enzymatic activity assays for TET2 using methylated single stranded DNA (ssDNA) as a substrate. The product was detected by a specific anti-5hmC antibody, and the activity of the enzyme was measured by immunocytochemistry using homogenous time resolved fluorescence (HTRF) technology. Again, we observed that R-2HG was a more potent inhibitor of TET2 than S-2HG, and that the maximum inhibition of TET2 with our assay was 97% for R-2HG and only 68% for S-2HG (Fig. 3F).

To gain a structural insight into the different inhibitory potencies of the 2HG metabolites, we investigated the conformation of KDM4C (PDB id 4XDO; resolution 1.97 Å) bound with aKG, S-2HG or R-2HG (Fig. 3G-I and Supp. Fig. 3E-F). Both S-2HG and R-2HG can bind to the active site of the enzyme, albeit in a distorted bidentate coordination when compared to aKG, since the keto carboxyl end of aKG is replaced by a hydroxyl group in S-2HG and R-2HG (Fig. 3J-L). The bidentate coordination of the active site of aKG-dependent enzymes is extremely sensitive to minimal changes (Domene et al., 2020), which explains why both S-2HG and R-2HG can act as inhibitors of KDM4C. Notably, we observed that R-2HG adopts a nearly identical orientation to a-KG in the catalytic core of KDM4C, in close proximity to Fe (II) (Fig. 3I,L and Supp. Fig. 3F). In contrast, S-2HG has to twist in order to find the right conformation in the catalytic core of KDM4C (Fig. 3H,K and Supp. Fig. 3E), which potentially explains why it is a less effective inhibitor. Collectively, these results highlight that S-2HG and R-2HG are not just isoforms of 2HG that indiscriminately target aKG-dependent enzymes, but that they have different potencies for different targets, and that this can result in different functions.

### OE-S-2HG treated mouse CD8+ T cells show enhanced tumour infiltration and increased effector molecule production

We next investigated the functional consequences of the observed differences between OE-S-2HG and OE-R-2HG treated CD8+ T cells in an *in vivo* murine adoptive cell transfer (ACT) *in vivo* model (Fig. 4A). Prior to adoptive T cell transfer, we determined their expression of CD44 (adhesion receptor), CD62L, CD25 (interleukin-2 receptor A), CTLA4 (receptor that acts as an immune checkpoint), ICOS (T cell co-stimulator) and Granzyme B (GzmB) (effector molecule) (Supp. Fig. 4A-B). Interestingly, even though mouse CD8+ T cells treated with OE-R-2HG do not lose CD62L expression, as their human counterparts do (Supp. Fig. 4A and Fig. 1E), we detected higher percentages of CD62L+/CD44+ (T_CM_) mouse CD8+ T cells when treated with OE-S-2HG compared to treatment with OE-R-2HG, or vehicle (Supp. Fig. 4A).

**Figure 4:**
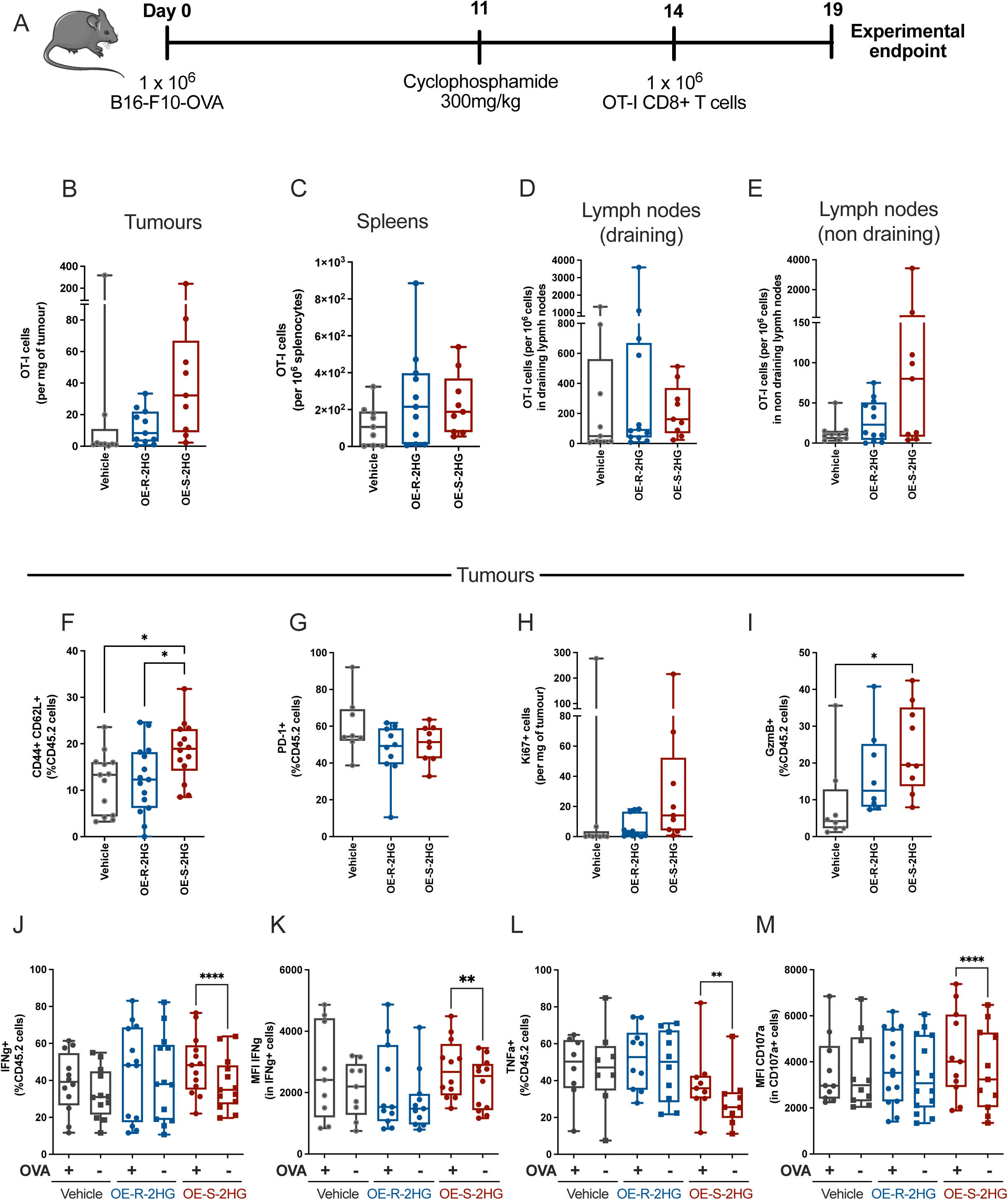
OE-S-2HG treated mouse CD8+ T cells are better at fine tuning effector molecule production. **(A)** Schematic representation of the adoptive cell therapy (ACT) *in vivo* model used. C57BL/6j (CD45.1+/CD45.2+) mice were injected subcutaneously with 1 million B16-F10 OVA-expressing tumour cells. On day 11, mice were lymphodepleted with 300 mg/kg cyclophosphamide. On Day 14, mice were intraperitoneally (i.p.) injected with PBS or 1 million CD45.2+ OT-I CD8+T cells, which were activated and treated with OE-S-2HG (0.4 mM), OE-R-2HG (0.4 mM) or vehicle (H_2_O) for 7 days *in vitro*. On day 19, the mice were sacrificed, and the tumours, spleens and draining / non-draining lymph nodes were harvested and analysed by flow cytometry. Endogenous and adoptive transferred populations were distinguished by the allelic variants of CD45. **(B)** Number of adoptively transferred OT-I cells per mg of tumour. The cell number was defined by counting beads. Median and min to max with all points shown (n= 9-11 mice per condition, two independent experiments). **(C)** Number of adoptively transferred OT-I cells per million splenocytes. The cell number was defined by counting beads. Median and min to max with all points shown (n= 9-11 mice per condition, two independent experiments). **(D-E)** Number of adoptively transferred OT-I cells (D) in draining and (E) non-draining lymph nodes. The cell number was defined by counting beads and the amount of the OT-I cells per million cells in lymph nodes was calculated. Median and min to max with all points shown (n= 9-11 mice per condition, two independent experiments). **(F)** Frequency of adoptively transferred OT-I cells (CD45.2+) expressing CD62L+/CD44+ markers infiltrated in the tumours. Median and min to max with all points shown (n= 14-15 mice per condition, three independent experiments). Unpaired two-tailed Student t test was used. **(G)** Frequency of adoptively transferred OT-I cells (CD45.2+) expressing PD-1 infiltrated in the tumours. Median and min to max with all points shown (n= 9-10 mice per condition, two independent experiments). **(H)** Number of adoptively transferred OT-I cells in the tumours positive for Ki67 per mg of tumour. The cell number was defined by counting beads. Median and min to max with all points shown (n= 9-11 mice per condition, two independent experiments). **(I)** Frequency of adoptively transferred OT-I cells (CD45.2+) expressing GzmB infiltrated in the tumours. Median and min to max with all points shown (n= 8-9 mice per condition, two independent experiments). Unpaired two-tailed Student t test was used. **(J-M)** Restimulation of OT-I tumour infiltrated lymphocytes *in vitro* with OVA_257-264_ (100 nM) peptide for 4 hrs. Brefaldin A and monensin were added at the last 2 hrs before flow cytometry analysis. (J) Frequency of adoptively transferred OT-I cells (CD45.2+) expressing IFNγ with (+OVA) or without (-OVA) restimulation. (K) IFNγ median fluorescent intensity (MFI) of adoptively transferred OT-I cells (CD45.2+, IFNγ+) cells with (+OVA) or without (-OVA) restimulation. (L) Frequency of adoptively transferred OT-I cells (CD45.2+) expressing TNFα with (+OVA) or without (-OVA) restimulation. (M) CD107a median fluorescent intensity (MFI) of adoptively transferred OT-I cells (CD45.2+, CD107a+) cells with (+OVA) or without (-OVA) restimulation. For panels J-M median and min to max with all points shown (n= 9-14 mice per condition, three independent experiments). Unpaired two-tailed Student t test was used between treatments and paired two-tailed Student t test was used for each treatment with (+OVA) or without (-OVA) restimulation.

B16-F10 OVA-bearing C57BL/6j mice treated with cyclophosphamide on day 11 received congenically marked OT-I CD8+ T cells at day 14 that were pre-treated with OE-S-2HG, OE-R-2HG, or vehicle (H_2_O) (Fig. 4A). Five days later, we determined the number of transferred OT-I T cells infiltrated in the tumours, spleens, and lymph nodes. The overall numbers of OT-I cells in the tumours were higher when T cells were pre-treated with OE-S-2HG, compared to vehicle or OE-R-2HG (Fig. 4B, Supp. Fig. 5A). Also, even though similar numbers of OT-I cells were found in spleens (Fig. 4C), mice that received OE-S-2HG treated OT-I cells had significantly bigger spleens than the mice that received vehicle treated OT-I cells (Supp. Fig. 5B). OE-S-2HG treated OT-I T cells were also found in increased numbers in non-draining lymph nodes, but not in draining lymph nodes, when compared to those found in animals given vehicle or OE-R-2HG treated cells (Fig. 4D-E).

Tumour-infiltrating OT-I T cells treated with OE-S-2HG had higher expression of CD44+/CD62L+ (T_CM_) than OE-R-2HG and vehicle treated cells (Fig. 4F). The activation marker PD-1 was unaltered between differentially treated OT-I tumour-infiltrating cells (Fig. 4G). We also checked the proliferation potential of the tumour infiltrated OT-I cells by staining for the proliferation marker Ki67. Even though the percentage of OT-I cells positive for Ki67 was similar between treatment groups (Supp. Fig. 5C), the number of Ki67+ cells was increased in tumours of mice that received OE-S-2HG treated OT-I cells compared to OE-R-2HG or vehicle treated T cells (Fig. 4H). We also found more GzmB-expressing cells in OE-S-2HG treated OT-I T cells, similarly to what was seen in our *in vitro* data (Fig. 4I and Supp. Fig. 4A-B).

It has been shown that tumour infiltrated lymphocytes (TILs) lose the capacity to react to T cell receptor (TCR) activation upon extraction from the tumour (Salerno et al., 2019). To investigate whether the treatment with the 2HG enantiomers could reverse this TCR block, we checked effector molecule production with or without *in vitro* restimulation with the cognate OVA_257-264_ peptide. We observed that OE-S-2HG treated OT-I cells significantly increased the percentage and the expression of the cytokine interferon-γ (IFNγ) specifically after restimulation (Fig. 4J-K), thus showing significantly decreased anergy. In addition, OE-S-2HG treated OT-I cells increased the percentage of positive cells, but not the expression levels, of the cytokine tumour necrosis factor-α (TNFα) after restimulation (Fig. 4L and Supp. Fig. 5D). We also checked the degranulation marker CD107a, and we found that its expression was significantly increased only in OE-S-2HG treated OT-I cells, again, specifically after restimulation (Fig. 4M and Supp. Fig. 5E). In contrast, OE-R-2HG and vehicle treated cells had similar cytokine responses before and after re-stimulation. The fact that a high percentage of OE-S-2HG treated OT-I cells were preserved in a T_CM_ (CD44+/CD62L+) phenotype within the tumour might explain the antigen-specific increase we observed when restimulating these cells. This would imply that the T_CM_ population could preserve these cells from exhaustion.

Finally, we analysed the adoptively transferred OT-I cells in the spleens, and performed *in vitro* restimulation with the OVA_257-264_ peptide. We observed a significant increase in the percentage of positive cells after antigen-restimulation for all of the conditions (vehicle, OE-S-2HG and OE-R-2HG) for IFNγ, interleukin 2 (IL-2) and CD107a, but a significant increase of TNFα+ after restimulation solely in the OE-S-2HG treated cells (Supp. Fig. 5F-I). Interestingly, in the unstimulated state (-OVA) OE-S-2HG treated OT-I splenocytes showed generally lower expression of cytokines compared to vehicle and OE-R-2HG, which, however, did not hinder the antigen-specific restimulation and cytokine expression of the OE-S-2HG treated cells (Supp. Fig. 5F-I). These data support the hypothesis that OE-S-2HG treated OT-I cells can preserve cytokine expression in the absence of antigen stimulation but are able to produce high amounts of effector molecules when needed.

### OE-S-2HG, but not OE-R-2HG treated mouse CD8+ T cells show increased anti-tumour activity

To determine the effect of both 2HG enantiomers on CD8+ T cell anti-tumour activity, we injected C57BL/6j mice with B16-F10 OVA-expressing tumour cells (Fig. 5A) and adoptively transferred OT-I CD8+ T cells that had been pre-treated with OE-S-2HG, OE-R-2HG or vehicle. At day 14 and day 21 after ACT we determined the number of OT-I T cells in circulation by flow cytometry (Fig. 5A). Irrespective of the day measured, we found significantly more OE-S-2HG treated OT-I cells in circulation compared to OE-R-2HG and vehicle treated OT-I T cells (Fig. 5B). Also, the percentage of CD62L+/CD44+ cells was higher for OE-S-2HG treated OT-I cells than for vehicle at all times measured, whereas the CD44+/CD62L+ population for OE-R-2HG treated OT-I cells was only higher at day 21 when compared to vehicle treated OT-I cells (Fig. 5C). We also checked the expression of CD127 (interleukin-7 receptor A) and found that OE-S-2HG treated OT-I cells had increased CD127 expression compared to vehicle at day 21 (Fig. 5D).

**Figure 5:**
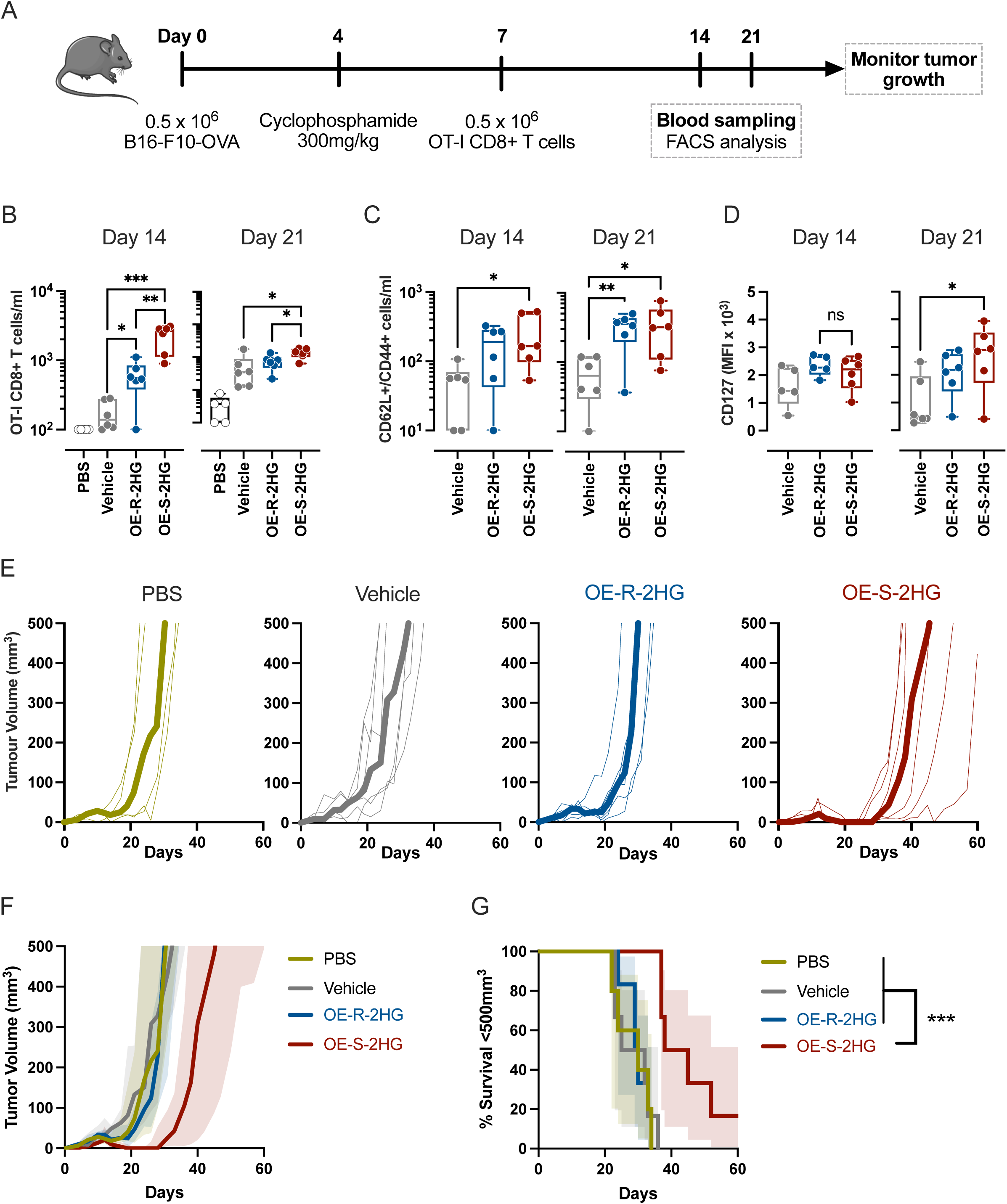
OE-S-2HG treated mouse CD8+ T cells show increase anti-tumour activity. **(A)** Schematic representation of the adoptive cell therapy (ACT) *in vivo* model used. C57BL/6j mice were injected subcutaneously with 0.5 million B16-F10 OVA-expressing tumour cells. On day 4, mice were lymphodepleted with 300 mg/kg cyclophosphamide. On Day 7, mice were intraperitoneally (i.p.) injected with PBS or 0.5 million CD45.1+ OT-I CD8+T cells, which were activated and treated with OE-S-2HG (0.4 mM), OE-R-2HG (0.4 mM) or vehicle (H_2_O) for 7 days *in vitro*. On day 14 and 21, peripheral blood was sampled and analysed by flow cytometry. Tumour growth was monitored every 2-3 days until day 60. **(B)** Frequency of adoptively transferred OT-I cells per millilitre of peripheral blood on day 14 (left) and day 21 (right). Median and min to max with all points shown of n= 5-6 mice per condition. Unpaired two-tailed Student t test was used. **(C)** Frequency of adoptively transferred OT-I cells positive for CD62L+/CD44+ markers per millilitre of peripheral blood on day 14 (left) and day 21 (right). Median and min to max with all points shown of n= 5-6 mice per condition. Unpaired two-tailed Student t test was used. **(D)** CD127 median fluorescent intensity (MFI) of adoptively transferred OT-I cells circulating in peripheral blood on day 14 (left) and day 21 (right). Median and min to max with all points shown of n= 5-6 mice per condition. Unpaired two-tailed Student t test was used. **(E)** Tumour growth for each condition. Thin lines represent tumour growth from individual mice and thick lines represent median tumour sizes for each group. **(F)** Combined data from (E). **(G)** Survival curves for tumour growth shown in (E-F). Threshold for survival was set at 500 mm^3^. Green line: PBS (n= 5 mice); grey line: vehicle (H_2_O) (n= 6 mice); blue line: OE-R-2HG (n= 6 mice); red line: OE-S-2HG (n= 6 mice). Representative of two independent experiments and log-rank (Mantel-Cox) test was used.

Finally, we measured the effect of 2HG enantiomers on T cells relative to the capacity of the cells to control tumour growth. Strikingly, OE-S-2HG treated OT-I cells showed superior anti-tumour activity compared to all other conditions tested (OE-R-2HG, vehicle and PBS - no T cell control), resulting in delayed tumour growth and increased survival (Fig. 5E-G). In contrast, OE-R-2HG OT-I cells showed no beneficial effects on anti-tumour activity compared to adoptive therapy of vehicle control treated T cells (Fig. 5E-G). This result can be explained by the increased number of tumour infiltrated lymphocytes (TILs) observed in the tumours (Fig. 4B), and by the fact that these TILs express higher levels of GzmB and can produce high amounts of cytokines after re-stimulation (Fig. 4I-M). Thus, the differences in gene expression, differentiation status and function observed in OE-S-2HG and OE-R-2HG treated CD8+ T cells result in altered anti-tumour activities.

## DISCUSSION

2HG is a by-product of the TCA cycle with important physiological roles. For example, S-2HG assists in adaptation to low oxygen levels (hypoxia) by regulating cellular redox state (Intlekofer et al., 2015; Xiao and Loscalzo, 2020). In addition, 2HG has significant roles in the fate decision of immune cells, and S-2HG levels are increased after CD8+ T cell receptor activation (Tyrakis et al., 2016).

Owing to its description as an oncometabolite, there is currently significant attention being paid to the role of 2HG in T cell immunology. This has resulted in somewhat contradictory findings. For example, Bunse, *et al.,* reported that non-cell permeable tumour-derived R-2HG can be taken up by human CD8+ T cells through the sodium-dependent dicarboxylate transporter 3 (SLC13A3), and that R-2HG accumulation in CD8 T cells leads to suppression of proliferation and anti-tumour activity (Bunse et al., 2018). Conversely, Bottcher, *et al.,* failed to detect impaired CD4+ or CD8+ T cell proliferation or increased cell death when cells were cultured with R-2HG. Rather, they found that R-2HG increased the frequency of regulatory T cells (Treg) and reduced the polarization of T helper 17 (Th17) cells (Bottcher et al., 2018). Other studies concluded that R-2HG accumulation in IDH mutant tumours inhibits both CD8+ T cell function and anti-tumour activity (Chuntova et al., 2022; Kadiyala et al., 2021; Wu et al., 2022; Zhang et al., 2018).

There are also contradictory data as far as S-2HG is concerned, although fewer studies have been conducted. Gupta, *et al.,* recently reported that pancreatic cancers accumulate S-2HG due to the promiscuous activity of LDHA, and by using an LDHA inhibitor they observed increased CD8+ T cell infiltration. Also, by using high concentrations of cell permeable S-2HG *in vitro*, the authors observed a reduction in CD8+ T cell migration and thus concluded that S-2HG suppresses anti-tumour activity (Gupta et al., 2021). In contrast to these results, our team has reported that exogenous treatment of mouse and human CD8+ T cells with S-2HG favours a memory phenotype and shows high anti-tumour activity in multiple adoptive T cell transfer (ACT) models (Foskolou et al., 2020; Tyrakis et al., 2016).

Here we demonstrate that in terms of T cell differentiation and function, 2HG is not one metabolite, and that its two enantiomers display qualitatively separable modes of action on CD8+ T cells and should thus be considered as two discrete entities. Specifically, treatment of human CD8+ T cells with cell permeable S-2HG or R-2HG differentially modulates CD8+ T cell proliferation and differentiation, which is reflected in their differential gene expression of transcription factors, CD molecules and secreted molecules. Importantly, OE-S-2HG treated CD8+ T cells expressed more memory-like markers on their surfaces and had a higher proliferative potential, a feature that is key for long term effective T cell responses *in vivo*. At the same time, OE-S-2HG treated cells retained expression of effector molecule transcripts. This was reflected also in our adoptive T cell experiments, where pre-treatment with OE-S-2HG increased the amount of functional CD8+ T cells present in the tumours, and showed that these cells had an increased cytotoxic potential.

We note here novel aspects of the molecular mechanisms of the divergent effects of the two 2HG variants. We found that OE-R-2HG treatment led to increased histone methylation in human CD8+ T cells, which implies potent inhibition of the aKG-dependent histone demethylases. Our *in vitro* enzymatic activity assay supported the notion that R-2HG is a more potent inhibitor than S-2HG of the KDM4C enzyme, which is in line with previous reports (Chowdhury et al., 2011). Interestingly, this is not the first time that S-2HG and R-2HG have been demonstrated to show different potencies in inhibiting aKG-dependent enzymes. Chowdhury *et al*., showed that S-2HG is a more potent inhibitor than R-2HG for the factor inhibiting HIF (FIH), prolyl hydroxylase domain 2 (PHD2) and alpha-ketoglutarate dependent dioxygenase homolog 2 (ALKBH2) enzymes, but they saw that R-2HG can inhibit the KDMs with greater potency (Chowdhury et al., 2011). Computational simulation assays showed that the backbone of S-2HG has to twist in order to adopt the pseudo pseudo-bidentate coordination to Fe(II) in the catalytic core of KDM4C, whereas R-2HG does not need to twist. This suggests that R-2HG can mimic aKG more closely, hence explaining the differential inhibitory activities of R-2HG and S-2HG.

We also observed that OE-R-2HG treatment decreased the levels of 5hmC DNA in human CD8+ T cells, likely due to inhibition of the aKG-dependent DNA demethylases (TET enzymes). Although R-2HG has been shown to act as a weak inhibitor for mouse TET1 and TET2 enzymes (Xu et al., 2011), our *in vitro* enzymatic activity assay with the human TET2 enzyme showed that R-2HG is a more potent inhibitor of this enzyme than S-2HG. These results indicate that S-2HG and R-2HG not only have different inhibitory potencies for different enzymes, but may also have distinct potencies for conserved enzymes in different species.

Finally, we investigated the functionality of OE-S-2HG and OE-R-2HG pre-treatment in adoptive cell transfer tumour models. Interestingly, OE-S-2HG treated OT-I cells preferentially migrated to tumour sites and were able to preserve their T_CM_ phenotype both within the tumour and in circulation. There were also more OE-S-2HG treated OT-I cells in the circulation compared to vehicle and OE-R-2HG treated cells, which can explain why we found more OT-I cells in the periphery (non-draining lymph nodes). More importantly, OE-S-2HG treated OT-I cells expressed higher levels of GzmB within tumours, and they were able to produce ample amounts of effector molecules specifically after antigen stimulation. This is an important characteristic of functional memory CD8+ T cells. The increased functionality of the OE-S-2HG pre-treated OT-I cells was demonstrated by their high anti-tumour activity and survival potential. Interestingly, although OE-R-2HG pre-treated OT-I cells did not show a beneficial anti-tumour effect, they also did not perform less well than the vehicle treated cells in our *in vivo* models.

In conclusion, OE-S-2HG pre-treatment before adoptive cell transfer increases CD8+ T cell fitness and enhances anti-tumour activity. Clearly, S-2HG and R-2HG are not merely enantiomeric forms but have distinct functions in CD8+ T cell biology.

## MATERIALS AND METHODS

### Cell culture and treatments

Peripheral blood mononuclear cells (PBMCs) were obtained from healthy donors from Cambridge Bioscience, National Health Service (NHS) Blood and Transplant (NHSBT: Addenbrooke’s Hospital, Cambridge, UK) or Sanquin (Amsterdam, NL). The study was performed according to the Declaration of Helsinki (seventh revision, 2013). Ethical approval was obtained from the East of England-Cambridge Central Research Ethics Committee (06/Q0108/281) and consent was obtained from all subjects. Written informed consent was obtained (Cambridge Bioscience, Cambridge, UK; NHSBT Cambridge, UK; Sanquin Research, Amsterdam, NL). Human CD8+ T cells were isolated either directly after blood donation (8-12 hrs after blood collection) or they were cryopreserved and used after cryopreservation. PBMCs were isolated through Ficoll-Paque PLUS density gradient separation (GE Healthcare). Cells were incubated in 21% oxygen, 5% carbon dioxide at 37 °C.

Human T cell isolation was performed with MACS Miltenyi kits (Naïve CD8+ T cells: 130-093-244 or Total CD8+ T cells: 130-096-495) following manufacturer’s instructions. CD8+ T cells were activated with aCD3/CD28 beads (1:1 beads-to-cell ratio) (11132D, Gibco). OE-S-2HG treatment and OE-R-2HG (H942595; H942596 Toronto Research Chemicals) started at day 0 and was at 0.4 mM concentration, otherwise stated. Every second day, fresh complete RPMI media (52400-025; Gibco) containing 10% FBS, 1% penicillin-streptomycin and IL-2 (30 U/ml) and the appropriate amount of OE-S-2HG, OE-R-2HG or vehicle was added. Cell number and viability were measured by ADAM-MC automated cell counter (NanoEnTek) or by CASY cell counter and analyser (BIOKE). Rate of cell division was calculated by dividing the cell number of day (X+2) by cell number of day X.

Mouse splenic OT-I CD8+ T cell isolation was performed with MACS Miltenyi kit (CD8+ T cells: 130-104-075) following manufacturer’s instructions. OT-I T cells were activated with either SIINFEKL (1000ng/mL) or by co-culturing them with pre-seeded OVA-expressing MEC.B7.SigOVA cells (van Stipdonk et al., 2003). The day after activation, the OT-I T cells were collected, washed, and treated with OE-S-2HG, OE-R-2HG or vehicle or 6-7 days. Cells were maintained in fresh complete RPMI media (52400-025; Gibco) containing 10% FCS, 1% penicillin-streptomycin, 55 μM β-mercaptoethanol and IL-2 (30 U/ml).

### Mice

C57BL/6J/Ly5.2, C57BL/6J/Ly5.1/Ly5.2 mice and C57BL/6J were bred in-house at the Netherlands Cancer Institute (NKI) or were purchased from Janvier Labs (for animal experiments performed in the Karolinska Institute, Sweden). Donor T cell receptor (TCR) transgenic mice (OT-I) mice were either crossed with mice bearing the CD45.1 congenic marker (002014, The Jackson Laboratory) for the tumour orthotopic models or with the CD45.2 congenic marker for the infiltration experiments. Experiments were performed in accordance with institutional and national guidelines and approved by the Experimental Animal Committee at the NKI and by the regional animal ethics committee of Northern Stockholm, Sweden under Ethical Permit number 5261-2020. All animals were housed in individually ventilated cage systems under specific-pathogen-free conditions. Both male and female mice were used at 8-12 weeks of age.

### Flow cytometry

Human CD8+ T cells FACS analysis was performed on day 12, otherwise stated. Mouse CD8+ T cells were analysed at day 7 of culture and OT-I adoptively transferred T cells were analysed after blood or organ/tumour harvesting. Cells were pelleted by centrifugation and stained with antibodies in FACS Buffer (5% FBS, 2 mM EDTA in PBS) at 4°C for 30–60 min. CCR7 staining was performed in media at 37°C. The stained cells were then washed with FACS buffer, pelleted, and re-suspended in 1x FACS-Fix (BD CellFIXTM) and kept at 4°C in the dark until processing. The samples were processed 2-3 days after fixation. For intracellular cytokine staining, cells were cultured in the presence of 1 μg/mL brefeldin A and monensin (00-4505-51, eBioscience) for 2 hr and were then fixed and permeabilized with Cytofix/Cytoperm kit (BD Biosciences) according to manufacturer’s protocol. For intracellular transcription factor and proliferation staining the Foxp3/Transcription Factor Staining (eBioscience) was used. On the day of analysis, the cells were resuspended in FACS Buffer containing counting beads (CountBrightTM Absolute Counting Beads). The number of cells in each sample was calculated according to the manufacturer’s instructions. Emission spectra “spillover” was corrected by compensation using compensation beads (01-1111-41; OneComp eBeads) mixed with each fluorescent probe. Flow cytometers used: BD LSR-Fortessa, AttuneX (Invitrogen) and BD FACSymphony. The flow data were analysed using FlowJo (BD Biosciences, version 10).

Human antibodies: Live/Dead (L34963, Invitrogen); CCR7 (3D12; BD Biosciences); CD28 (CD28.2; Biolegend); CD45RO (UCHL1; Biolegend); CD62L (DREG-56; Biolegend); CD8 (HIT8a; Biolegend); PD-1 (EH12.2H7; Biolegend); TOX (TXRX10; Fisher Scientific); 5hmC (10013602; Active Motif). Mouse antibodies: Live/Dead Near-IR (L10119, Invitrogen); CD107a (ID4B, eBiosciences); CD127 (A7R34, Biolegend); CD25 (PC61.5, eBiosciences); CD44 (IM7, Biolegend); CD45.1 (A20, Biolegend); CD45.2 (104, Biolegend); CD62L (MEL-14, Biolegend); CD8a (53-6.7, Biolegend); CTLA-4 (UC10-4B9, Biolegend); FcR Block (120-000-826, MACS Miltenyi); GzmB (AD2 and GB11, Biolegend and BD); Ki67 (SolA15 and B56, eBiosciences and BD); ICOS (C398.4A, Biolegend); IFN-γ (XMG1.2, Biolegend); IL-2 (JES6-5H4, eBiosciences); PD-1 (J43, eBiosciences); TNF-α (MP6-XT22, Biolegend).

### Western Blots and 5hmC staining

For western blots, human naïve CD8+ T cells were isolated from healthy donors and treated from day 0 to day 12 with OE-S-2HG (0.4 mM), OE-R-2HG (0.4 mM) or vehicle. The cells were activated with aCD3/CD28 beads for 4 days as explained above. At day 12, cells were collected, washed, lysed and the histones were extracted by using the Histone Extraction Kit (ab113476, Abcam) the manufacturer’s instructions. Protein quantification was performed with the BCA protein assay kit (ab207003, Abcam). Proteins were separated by SDS-PAGE and transferred to PVDF membranes. Membranes were then blocked in 5% milk prepared in PBS with 0.05% Tween 20, incubated with primary antibodies overnight at 4 ⁰C and HRP-conjugated secondary antibodies (HAF008 and HAF007, R&D) for 1 hr at room temperature the next day. Following ECL exposure (GERPN2106, Sigma), membranes were imaged using an iBrightCL1000 (Thermo Fisher). The following primary antibodies were used for western blots at a concentration (1:1000) from Cell Signalling H3K9Ac (9649); H3K9me2 (4658); H3K9me3 (13969); H3K27Ac (8173); H3K27me3 (9733); H3 (4499).

For 5hmC staining, total CD8+ T cells were isolated from healthy donors and treated from day 0 to day 7 with OE-S-2HG (0.4 mM), OE-R-2HG (0.4 mM) or vehicle. The cells were activated with aCD3/CD28 beads for 4 days as explained above. At day 7, cells were collected, washed, and stained with Live/Dead (L34963, Invitrogen). The cells were then fixed with the True-Nuclear Transcription Factor kit (Biolegend). After fixation and permeabilization, the cells were incubated with 4M HCL for 10 min at room temperature. The cells were then thoroughly washed and incubated in blocking buffer (0.1% PBS-Triton, 5% FBS) for 30min at 4°C. The cells were then incubated with primary anti-5hmC (10013602; Active Motif) overnight at 4°C and the day after with secondary antibody for 1 hr at room temperature. Flow cytometry was then performed as explained above.

### RNA-Seq analysis and GSEA

Naïve CD8 T cells activated and treated with OE-S-2HG (0.4 mM), OE-R-2HG (0.4 mM) or vehicle and cells were collected on day 5 or day 12. The cells were lysed on RLT Buffer containing (1:100) β-Mercaptoethanol. RNA-Seq libraries and data analysis was performed by Active Motif. DESeq2 was used for differential analysis statistics. For read mapping, the RNA-sequencing generated 42-nt sequence reads using Illumina NextSeq 500. The reads were mapped to the genome using the STAR algorithm with default settings. Only read pairs that have both ends aligned were counted. Read pairs that have their two ends mapping to different chromosomes or mapping to same chromosome but on different strands were discarded. Also, at least 25 bp overlapping bases in a fragment for read assignment were required. GSEA analysis was applied from the entire list of genes that compose the RNA-Seq expression matrix. The input used a gene list ranked by shrunkenLog2FC obtained from DESeq2, we performed GSEA with default setting to determine whether members of a priori defined gene set based on biological knowledge (e.g. genes sharing the same GO category) tend to occur toward the top (or bottom) of the gene list. GSEAPreranked tool developed by Broad Institute: (http://software.broadinstitute.org/gsea/index.jsp).

The OE-R-2HG upregulated significant genes were used for enrichment analysis with the EnrichR tool (Kuleshov et al., 2016) and the ENCODE transcription factor ChIP-Seq library. The Human Protein Atlas (proteinatlas.org) and Monaco (Monaco et al., 2019) dataset was used to check the expression of specific genes in different immune cell types.

### *In vitro* enzymatic activity assays

For enzymatic activity of the KDM4C enzyme, the human KDM4C enzyme (8 nM, BPS Bioscience) was incubated with the substrate of H3(1-21) lysine 9 tri-methylated biotinylated peptide (30 nM, Anaspec), in the presence of aKG (10 μM, Sigma), ammonium iron(II) sulfate hexahydrate (5 μM, Sigma), assay buffer (50 mM HEPES pH 7.0, 0.01% Tween 20, 1mM ascorbic acid and 0.01% BSA). S-2HG (Sigma) and R-2HG (Sigma) were used in 3-fold serial dilution and the maximum concentration used was 5 mM. The pre-incubation time of the inhibitors with the KDM4C mixture was 30 min at room temperature. The reaction step was at room temperature for 210 min. The product was detected by using an anti-histone H3K9 Me2-Eu(K) antibody (Cisbio) and XL665-conjugated Streptavidin (SA-XL665, Cisbio). The detection of the homogenous time resolved fluorescence (HTRF) signal was proportional to the concentration of demethylated H3(1-21) peptide. The assays were performed in technical duplicates in a 384-well plate. The inhibitor analysis was performed within the linear range of catalysis. For enzymatic activity of the TET2 enzyme, the human TET2 enzyme (2 nM, BPS Bioscience) was incubated with the substrate of ssDNA (ssBiotin 26nt Me-C Oligo 30 nM, Genscript), in the presence of aKG (115 μM, Sigma), ammonium iron(II) sulfate hexahydrate (10 μM, Sigma), in assay buffer (50 mM HEPES pH 7.0, 100 mM NaCl, 0.01% Pluronic F-127, 1mM TCEP, 2mM ascorbic acid, 0.2 mg/ml BSA and 1000U/ml Catalase). S-2HG (Sigma) and R-2HG (Sigma) were used in 3-fold serial dilution and the maximum concentration used was 5 mM. The pre-incubation time of the inhibitors with the TET2 mixture was 30 min at room temperature. The reaction step was at room temperature for 90 min. The product was detected by using an anti-5-Hydroxymethylcytosine antibody (5 nM, Active Motif), Eu-Protein A (5 nM, Cisbio), Streptavidin-Alexa Fluor 647 (6.25 nM, Life Technologies) and 10 mM EDTA (Sigma). For the standard curve, the ssBiotin 26nt HydMe-C Oligo (Genscript) was used. The assays were performed in technical duplicates in a 384-well plate. The inhibitor analysis was performed within the linear range of catalysis. The percentage of inhibition was calculated with the following formula: Inhibition%=(1- (signal value per well-Average Low control)/(Average High control-Average Low control))*100. The data were fitted by Prism Graphpad with four parameters equation via “log(inhibitor) vs. response -- Variable slope” model.

### Simulations

#### System setup

The atomic structure of the KDM4C protein (PDB id 4XDO; resolution 1.97 Å) (Wigle et al., 2015) in complex with aKG (KDM4C:aKG:Fe(II)) was used as a starting point for Molecular Dynamics (MD) simulations. Three systems were prepared: KDM4C:aKG:Fe(II), KDM4C:2HG-(*S*):Fe(II) and KDM4C:2HG-(*R*):Fe(II). Calculations were done with NAMD 2.9, (Phillips et al., 2005) using the CHARMM 36 protein force-field (Brooks et al., 2009) together with the TIP3P water model (Jorgensen et al., 1983). 2OG, 2HG-(*S*), and 2HG-(*R*) parameters were obtained in CHARMM from Paramchem (Vanommeslaeghe and MacKerell, 2012). Fe(II) Lennard Jones parameters were taken from the CHARMM force field. Default ionization states were used for the protein on the basis of PropKa calculations (Rostkowski et al., 2011).

#### Simulation details

The particle mesh Ewald (PME) algorithm was used for the evaluation of electrostatic interactions beyond 12 Å, with a PME grid spacing of 1 Å, and NAMD defaults for spline and κ values (Darden et al., 1993). A cutoff at 12 Å was applied to nonbonded forces. Both electrostatics and van der Waals forces were smoothly switched off between the cutfoff distance of 12 Å and the switching distance of 10 Å using the default NAMD switching function. A Verlet neighbour list (Verlet, 1967) with pairlist distance of 14 Å was used only to evaluate nonbonded neighboring forces within the pairlist distance. The lengths of covalent bonds involving hydrogen atoms were constrained by the SETTLE algorithm (Miyamoto and Kollman, 1992) to be able to use a 2 fs timestep. The multi time step algorithm Verlet-I/r-RESPA (Tuckerman et al., 1992) was used to integrate the equations of motion. Nonbonded short-range forces were computed for each time step, while long-range electrostatic forces were updated every 2 timesteps. The pressure was kept at 1.026 bar (1 atm) by the Nosé-Hoover Langevin piston (Nosé, 1984a; Nosé, 1984b), with a damping time constant of 50 fs and a period of 100 fs. The temperature was maintained at 300 K by coupling the system to a Langevin thermostat (Langevin, 1908), with a damping coefficient of 5 ps^-1^. A 150 mM background ionic concentration of NaCl was utilized to achieve system neutrality. The total system size was 94,691 atoms, comprised of 8,264 waters, 84 Na^+^ ions and 88 Cl^-^ ions.

After 1,000 steps of Conjugate-Gradient minimization with restraints on the protein and co-factor, and 10 ns of simulation with backbone restraints and restraints on the co-factor, a 50 ns production run in the NPT ensemble was carried out for each system.

### Animal studies

For infiltration experiments, 8-12-week-old C57BL/6J/Ly5.1/Ly5.2 mice were injected subcutaneously with 1 x 10^6^ B16-F10-OVA cells and conditioned 11 days later with peritoneal injection of 300 mg/kg cyclophosphamide (Sigma, #C0768). On day 14, 1 x 10^6^ OT-I Ly5.2 CD8+ T cells were peritoneally injected (the OT-I cells were previously activated and treated with 0.4 mM OE-S-2HG, OE-R-2HG or vehicle *in vitro* for 7 days). Animals were assigned randomly to each experimental group. On day 19, tumours, spleens, and lymph nodes (draining and non-draining) were harvested. The excised tumours were cut into small pieces and digested with 100 μg/ml DNase I (Roche) and 200 U/ml Collagenase (Worthington) at 37°C for 30 min. The spleens and lymph nodes were smashed over a 40 μm filter. Cells were counted, and where indicated, they were re-stimulated for 4 hr with 100 nM OVA_257–264_ peptide and brefeldin A and monensin were added for the last 2 hr of activation. The tumour single-cell suspensions were stained with fluorochrome-labelled antibodies and analysed by flow cytometry.

For tumour growth experiments, 8-15-weeks-old female C57BL/6J/Ly5.2 were inoculated subcutaneously with 0.5 x 10^6^ B16-F10-OVA and conditioned 4 days later with peritoneal injection of 300 mg/kg cyclophosphamide (Sigma). On day 7, 0.5 x 10^6^ OT-I Ly5.1 CD8+ T cells were peritoneally injected (the OT-I cells were previously activated and treated with 0.4 mM OE-S-2HG, OE-R-2HG or vehicle *in vitro* for 7 days). Animals were assigned randomly to each experimental group. Tumour volume was measured every 2-3 days with electronic callipers until day 60. Peripheral blood was collected from the tail vein at days 14 and 21 and analysed by flow cytometry. Tumour volume was calculated using the formula a×b×b/2 where a is the length and b is the width of the tumour. Mice were sacrificed when the tumours reached a size of 500 mm^3^.

### Statistical Analysis

Statistical analysis was performed with Prism-9 software (Graph-Pad). Statistical significance was set at p<0.05 and the statistical tests used are stated in figure legends.

## ACKNOWLEDGMENTS

We thank Darren Cawkill, Mark Hoogenboezem, Ken GC Smith, and Pedro Velica. The Cambridge NIHR BRC Cell Phenotyping Hub and the flow cytometry facility from the School of the Biological Sciences. The animal caretakers of the NKI and the Sanquin FACS facility. This work was supported by Apollo Therapeutics and Wellcome Trust Principal Fellowship Award to RSJ; the Evelyn Trust Cambridge (Patrick Sisson’s Research Fellowship) and the Karolinska Institutet (Jonas Söderquists Fellowship) awarded to IPF; the Foundation for Science and Technology scholarship (SFRH/BD/115612/2016) awarded to PPC.

## AUTHORSHIP CONTRIBUTIONS

IPF conceived, designed, performed, and interpreted the majority of the experiments and wrote the manuscript. PPC conducted and analysed experiments (including western blot and *in vivo* experiments) and reviewed the manuscript. EM conducted and analysed experiments (including 5hmC staining) and reviewed the manuscript. BPN and DB analysed the RNA-Seq data and reviewed the manuscript. AG and NDZ conducted *in vivo* experiments and reviewed the manuscript. CJ conducted the simulation assays and reviewed the manuscript. LB, DN, PT and AS conducted and analysed experiments and/or reviewed the manuscript. MCW supervised *in vivo* experiments and reviewed the manuscript. RSJ designed the study, wrote the manuscript, and supervised the project.

---

**Supp. Figure 1:**
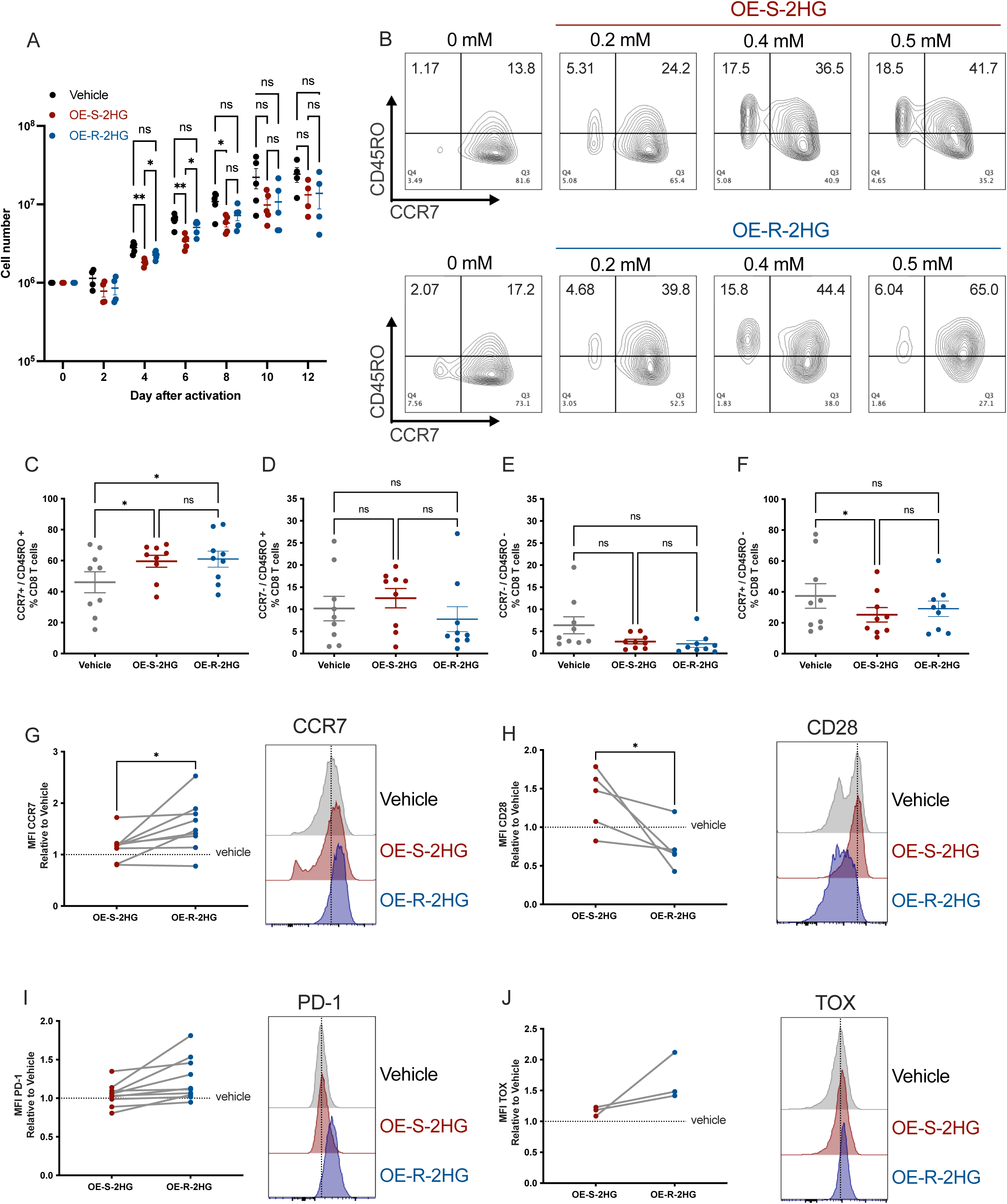
Expression of surface markers in human CD8+ T cells treated with OE-S-2HG or OE-R-2HG. **(A)** Cell number of CD8+ T cells treated with OE-S-2HG (0.4 mM), OE-R-2HG (0.4 mM) or vehicle (H_2_O) for the indicated days as determined by an automated cell counter. Data are represented in log10 as mean ± SEM. Mixed-effects analysis with Tukey’s multiple comparison test was used. **(B)** Flow cytometry plots of CD8+ T cells showing surface expression of CCR7 and CD45RO. Cells were treated with vehicle (H_2_O) or increasing concentrations of OE-S-2HG or OE-R-2HG and analysed at day 12 by flow cytometry. Representative plots of n= 3 is shown. **(C-F)** Cells were treated with OE-S-2HG (0.4 mM), OE-R-2HG (0.4 mM) or vehicle (H_2_O) and the proportion of (C) CCR7+/CD45RO+; (D) CCR7-/CD45RO+; (E) CCR7-/CD45RO-; (F) CCR7+/CD45RO-cells is shown (%CD8+ T cells). Cells were analysed at day 12/13 by flow cytometry. Each data point represents a donor (n= 9; from 5 independent experiments). Data are represented as mean ± SEM. RM one-way ANOVA with Tukey’s multiple comparison test was used. **(G)** Fold change of median fluorescence intensity (MFI) of CCR7 for CD8+ T cells treated with OE-S-2HG (0.4 mM) or OE-R-2HG (0.4 mM) relative to vehicle (H_2_O) on the left, and representative histogram flow cytometry plots on the right. Cells were analysed at day 12. Each data point represents a donor (n= 9; from 6 independent experiments). Unpaired two-tailed Student t test was used. **(H)** Fold change of median fluorescence intensity (MFI) of CD28 for CD8+ T cells treated with OE-S-2HG (0.4 mM) or OE-R-2HG (0.4 mM) relative to vehicle (H_2_O) on the left, and representative histogram flow cytometry plots on the right. Cells were analysed at day 12/13. Each data point represents a donor (n= 5; from 3 independent experiments). Unpaired two-tailed Student t test was used. **(I)** Fold change of median fluorescence intensity (MFI) of PD-1 for CD8+ T cells treated with OE-S-2HG (0.4 mM) or OE-R-2HG (0.4 mM) relative to vehicle (H_2_O) on the left, and representative histogram flow cytometry plots on the right. Cells were analysed at day 12. Each data point represents a donor (n= 10; from 6 independent experiments). Unpaired two-tailed Student t test was used. **(J)** Fold change of median fluorescence intensity (MFI) of TOX for CD8+ T cells treated with OE-S-2HG (0.4 mM) or OE-R-2HG (0.4 mM) relative to vehicle (H_2_O) on the left, and representative histogram flow cytometry plots on the right. Cells were analysed at day 15. Each data point represents a donor (n= 3). Unpaired two-tailed Student t test was used. For all panels naïve CD8+ T cells were isolated and activated with CD3/CD28 beads and cultured with IL2 (30 U/mL) in the presence of OE-S-2HG (0.4 mM), OE-R-2HG (0.4 mM) or vehicle (H_2_O) from day 0 to 12, unless otherwise stated.

**Supp. Figure 2:**
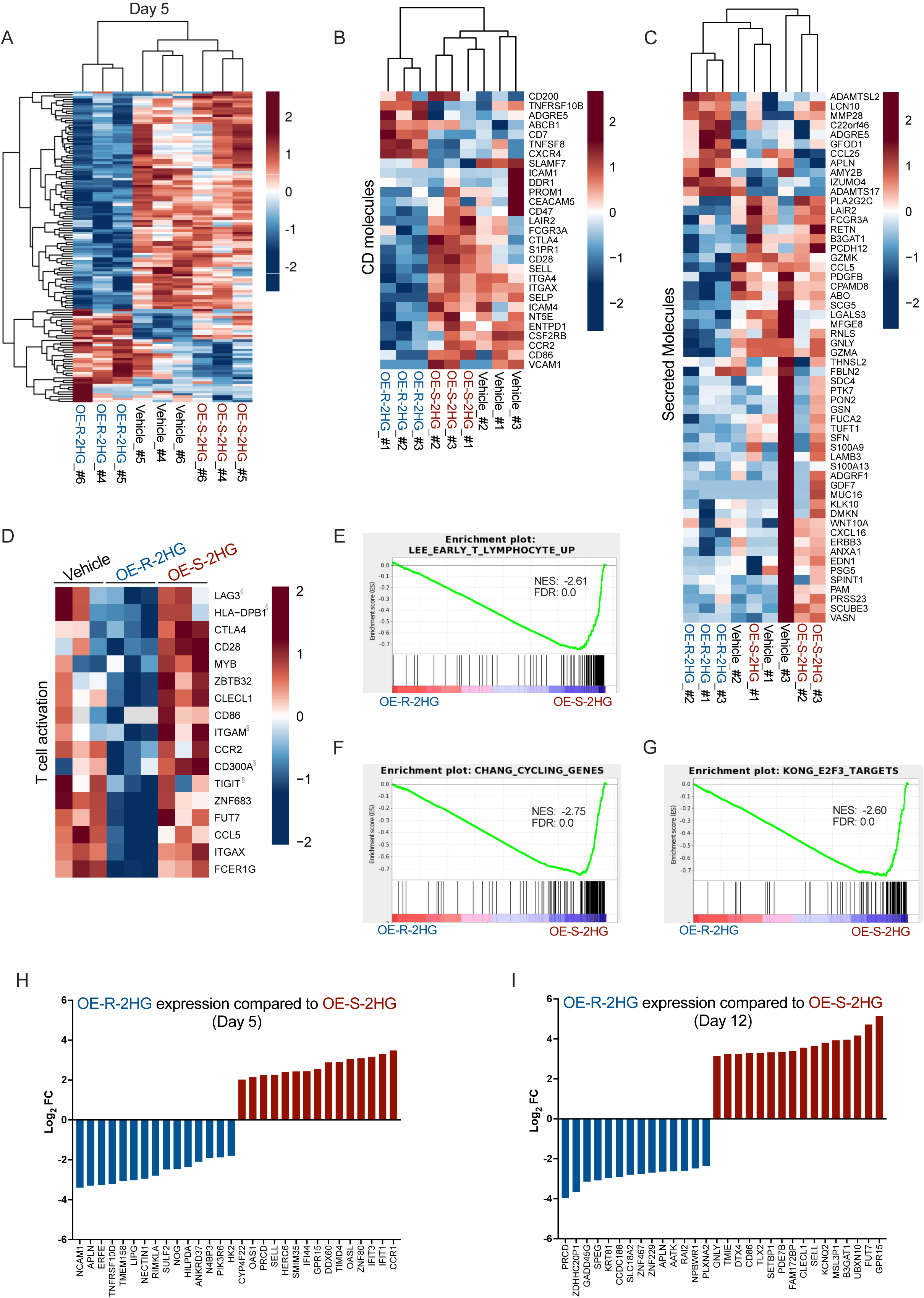
RNA-Seq analysis of OE-S-2HG and OE-R-2HG treated human CD8+ T cells. Naïve CD8+ T cells were isolated from 6 individual donors over 3 independent experiments, activated and treated with OE-S-2HG (0.4 mM), OE-R-2HG (0.4 mM) or vehicle (H_2_O). The cells were collected either on day 5 (3 donors) or on day 12 (3 donors) and RNA-Seq analysis followed. **(A)** Heatmap of hierarchically clustered genes in CD8+ T cells treated with OE-S-2HG (0.4 mM), OE-R-2HG (0.4 mM) or vehicle (H_2_O) at day 5 of culture. **(B-C)** Heatmaps of hierarchically clustered genes in CD8+ T cells treated with OE-S-2HG (0.4 mM), OE-R-2HG (0.4 mM) or vehicle (H_2_O) at day 12 of culture. Statistically significant differentially expressed hits of (B) CD molecules and (C) secreted molecules are shown. **(D)** Heatmap of standardized gene expression (Z score) in treated CD8+ T cells of genes involved in T cell activation. The gene set was obtained from ToppGene. Red and blue colours indicate increased and decreased expression respectively. Genes marked with ‘§’ were not statistically significant hits. Samples from day 12 of treatment are shown. **(E-G)** Gene set enrichment analysis (GSEA) of treated CD8+ T cells at day 12 of culture for (E) Lee early T lymphocytes up; (F) Chang cycling genes and (G) Kong E2F3 targets. Net enrichment score (NES) values and false discovery rate (FDR) are shown. **(H-I)** Plots show the top 30 most differentially expressed genes for (H) day 5 and (I) day 12 of CD8+ T cells treated with OE-S-2HG (0.4 mM) or OE-R-2HG (0.4 mM).

**Supp. Figure 3:**
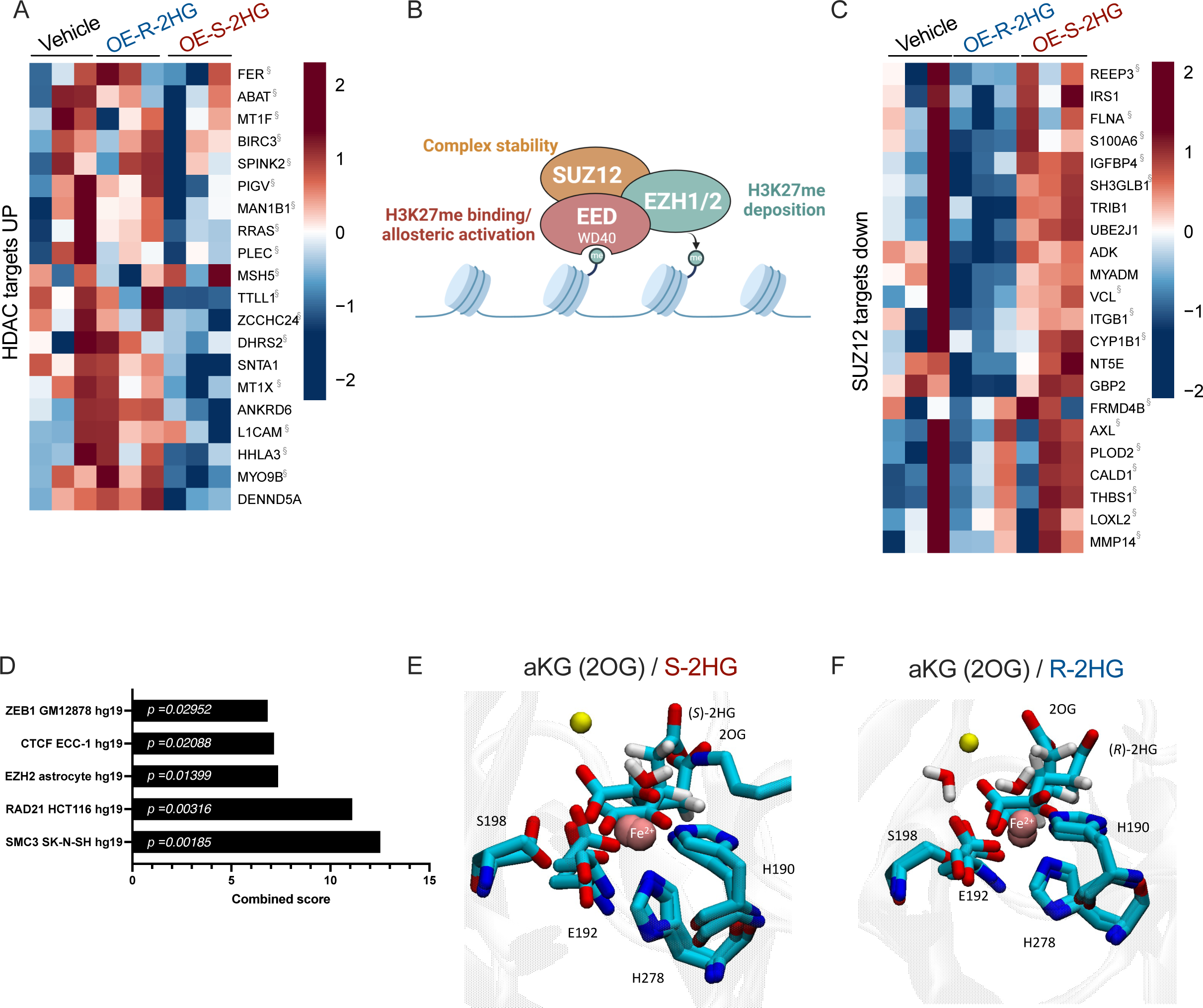
Mechanistic insight of OE-S-2HG and OE-R-2HG treatment in CD8+ T cells. **(A)** RNA-Seq was performed as described in figure 2. Heatmap of standardized gene expression (Z score) in treated CD8+ T cells for HDAC gene targets (upregulated). The gene set was obtained from ToppGene. Red and blue colours indicate increased and decreased expression respectively. Genes marked with ‘§’ were not statistically significant hits. Samples from day 12 of treatment are shown. **(B)** Schematic representation of the PRC2 complex is shown. **(C)** RNA-Seq was performed as described in figure 2. Heatmap of standardized gene expression (Z score) in treated CD8+ T cells for SUZ12 targets down (downregulated). The gene set was obtained from ToppGene. Red and blue colours indicate increased and decreased expression respectively. Genes marked with ‘§’ were not statistically significant hits. Samples from day 12 of treatment are shown. **(D)** OE-R-2HG upregulated significant hits were used for enrichment analysis with the EnrichR tool (Kuleshov et al., 2016) using the the ENCODE TF ChIP-seq 2015 data set. The combined scores of significant targets are shown. **(E)** Overlay images of the catalytic site of KDM4C protein (PDB id 4XDO; resolution 1.97 Å) in complex with aKG (KDM4C:2OG:Fe(II)) and S-2HG (KDM4C:2HG-(S):Fe(II)). **(F)** Overlay images of the catalytic site of KDM4C protein (PDB id 4XDO; resolution 1.97 Å) in complex with aKG (KDM4C:2OG:Fe(II)) and R-2HG (KDM4C:2HG-(R):Fe(II)).

**Supp. Figure 4:**
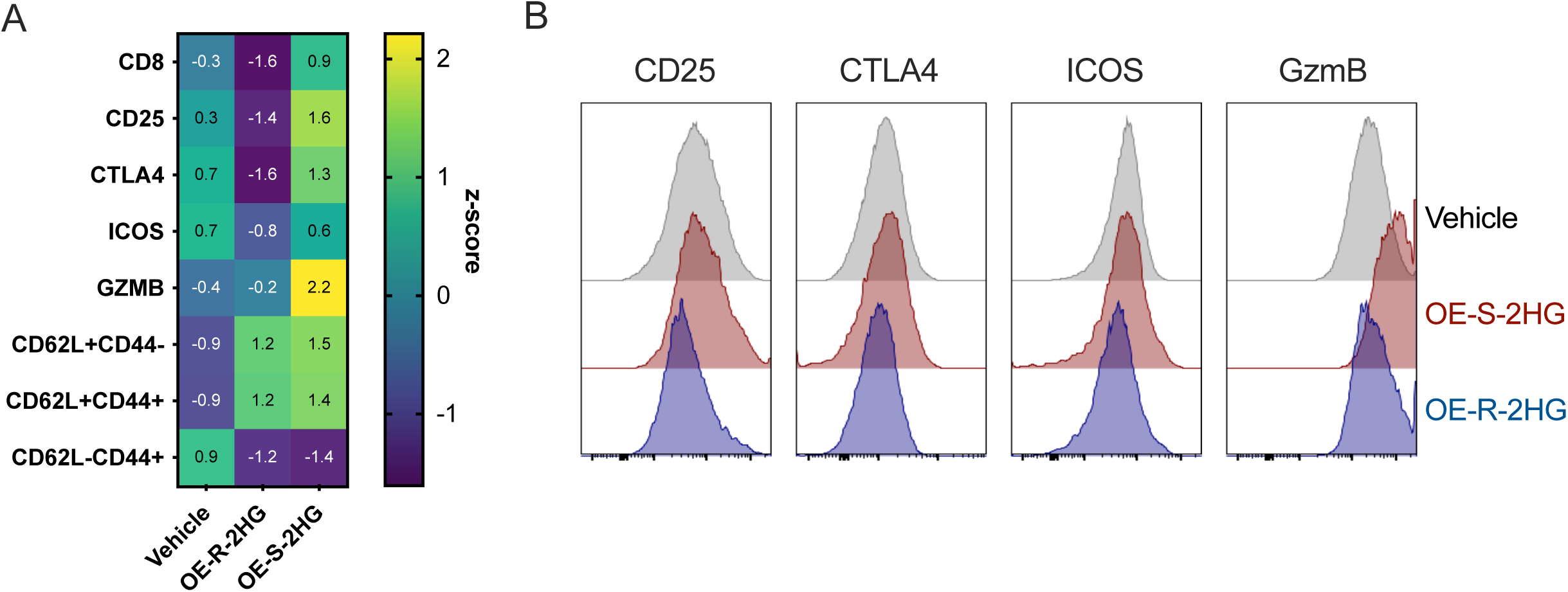
Analysis of OE-S-2HG and OE-R-2HG treated mouse CD8+ T *in vitro*. **(A)** Heatmap of standardized expression (Z score) for specific markers or percentage of specific populations in OT-I CD8+ T cells treated with vehicle (H_2_O), OE-R-2HG (0.4 mM) or OE-S-2HG (0.4 mM) for 7 days *in vitro*. **(B)** Representative histogram flow cytometry plots of the markers tested in (A).

**Supp. Figure 5:**
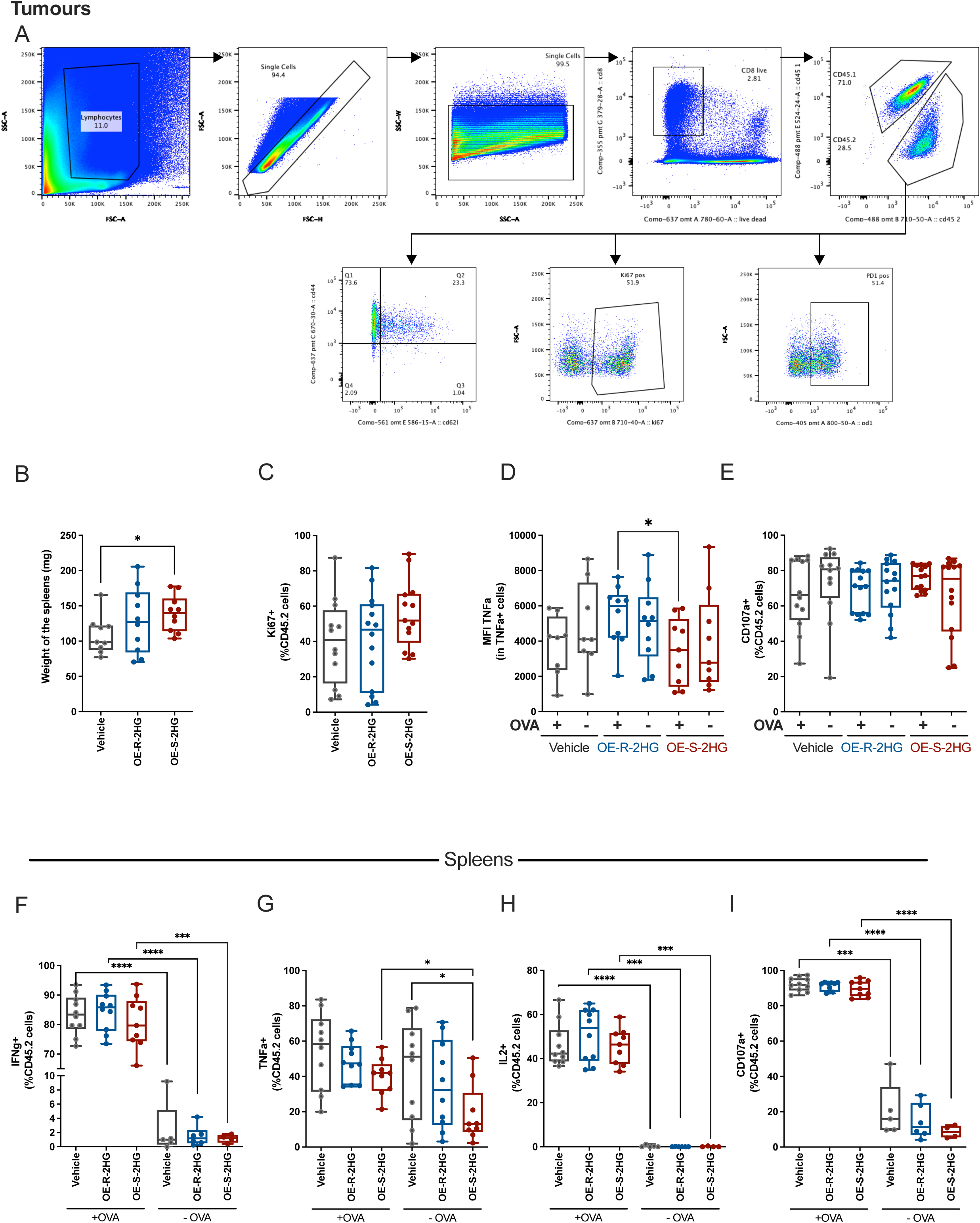
OE-S-2HG and OE-R-2HG treated mouse CD8+ T cells in adoptive cell transfer model. **(A)** Gating strategy of the OT-I tumour infiltrated lymphocytes. The host CD8+ T cells were CD45.1+CD45.2+ positive and the adoptive transferred OT-I cells were CD45.2+. **(B)** Weights of spleens from mice treated with OE-S-2HG, OE-R-2HG or vehicle OT-I cells. **(C)** Frequency of adoptively transferred OT-I cells (CD45.2+) expressing Ki67+ infiltrated in the tumours. Median and min to max with all points shown (n= 12-14 mice per condition, three independent experiments). **(D-E)** Restimulation of OT-I tumour infiltrated lymphocytes *in vitro* with OVA_257-264_ (100 nM) peptide for 4 hrs. Brefaldin A and monensin were added at the last 2 hrs before flow cytometry analysis. (D) TNFα median fluorescent intensity (MFI) of adoptively transferred OT-I cells (CD45.2+, TNFα+) cells with (+OVA) or without (-OVA) restimulation. (E) Frequency of adoptively transferred OT-I cells (CD45.2+) expressing CD107a with (+OVA) or without (-OVA) restimulation. **(F-I)** Restimulation of OT-I spleen infiltrated lymphocytes *in vitro* with OVA_257-264_ (100 nM) peptide for 4 hrs. Brefaldin A and monensin were added at the last 2 hrs before flow cytometry analysis. Frequency of adoptively transferred OT-I cells (CD45.2+) expressing (F) IFNγ, (G) TNFα, (H) IL2, (I) CD107a with (+OVA) or without (-OVA) restimulation. For panels D-E median and min to max with all points shown (n= 9-14 mice per condition, three independent experiments). For panels F-I median and min to max with all points shown (n= 5-10 mice per condition, two independent experiments). Unpaired two-tailed Student t test was used between treatments and paired two-tailed Student t test was used for each treatment with (+OVA) or without (-OVA) restimulation.

